# The ecology of the *Drosophila*-yeast mutualism in wineries

**DOI:** 10.1101/273839

**Authors:** Allison S. Quan, Michael B. Eisen

## Abstract

The fruit fly, *Drosophila melanogaster,* is preferentially found on fermenting fruits. The yeasts that dominate the microbial communities of these substrates are the primary food source for developing *D. melanogaster* larvae, and adult flies manifest a strong olfactory system-mediated attraction for the volatile compounds produced by these yeasts during fermentation. Although most work on this interaction has focused on the standard laboratory yeast *Saccharomyces cerevisiae*, a wide variety of other yeasts naturally ferment fallen fruit. Here we address the open question of whether *D. melanogaster* preferentially associates with distinct yeasts in different, closely-related environments. We characterized the spatial and temporal dynamics of *Drosophila*-associated fungi in Northern California wineries that use organic grapes and natural fermentation using high-throughput, short-amplicon sequencing. We found that there is nonrandom structure in the fungal communities that are vectored by flies both between and within vineyards. Within wineries, the fungal communities associated with flies in cellars, fermentation tanks, and pomace piles are distinguished by varying abundances of a small number of yeast species. To investigate the origins of this structure, we assayed *Drosophila* attraction to, oviposition on, larval development in, and longevity when consuming the yeasts that distinguish vineyard microhabitats from each other. We found that wild fly lines did not respond differentially to the yeast species that distinguish winery habitats in habitat specific manner. Instead, this subset of yeast shares traits that make them attractive to and ensure their close association with *Drosophila*.

## Introduction

All animals interact with microbes, and it is increasingly clear that the collection of microbes with which an animal interacts can have a dramatic impact on its physiology, behavior, and other phenotypes [1–5]. Some of the microbes associated with animals in the wild are highly specific and acquired through dedicated mechanisms that ensure the robust maintenance of their interaction [6–8]. Other associations, however, are more contingent, and involve microbes acquired as the animal navigates a microbe rich environment. While this latter class has received less attention, studying the contingent microbiome of wild animals can reveal important details of natural history, ecology and behavior.

The microbiome of the fruit fly, *Drosophila melanogaster,* represents an interesting mix of obligate and contingent microbial associations [9]. In nature, *D. melanogaster* is found on or near fermenting substrates, on which they preferentially oviposit, as *D. melanogaster* larvae (and indeed those of most *Drosophila* species) feed on microbes, particularly yeasts [10,11]. Yeasts benefit from visits by adult flies, who vector them from site to site, enabling their dispersal and colonization of new substrates [12,13]. This association is mediated by a strong, olfactory-based attraction of adult *D. melanogaster* to the volatile compounds produced during yeast fermentation [14–16].

A growing body of work has investigated the interaction between *D. melanogaster* and the brewer’s yeast, *Saccharomyces cerevisiae,* in the laboratory. Adult fruit flies prefer substrates inoculated with yeast over any other sterile substrate [15], and under laboratory conditions, *D. melanogaster* can discriminate between and prefers some strains of *S. cerevisiae* over others based on volatile profile alone [17–19]. While the interaction between flies and yeasts is clear, the specificity of this interaction in nature has been poorly studied.

Both *D. melanogaster* and *S. cerevisiae* are found in abundance in wineries, a habitat more natural than a controlled laboratory but more accessible than a completely wild ecosystem [20]. A variety of non-*Saccharomyces* yeasts are observed during spontaneous fermentation, a winemaking practice in which only the yeasts found on the grapes at the time of harvest, and those introduced naturally or incidentally after harvest, are used for fermentation [21,22]. However, little is understood about the movement of non-*Saccharomyces* yeasts in vineyards, although insects are acknowledged as potentially important vectors [23–27]. Given that drosophilids are closely associated with yeast throughout their entire lifecycle, flies are a likely candidate for vineyard and winery yeast dispersal [13,28,29]. However, the yeasts associated with vineyard and winery *Drosophila* have yet to be thoroughly characterized.

In a broader context, investigating the degree of specificity of the fly-yeast mutualism in nature can help reveal both the constraints and plasticity of natural mutualisms. While several studies have characterized the yeasts vectored by *Drosophila* in vineyards and wineries using culture-based methods [20,30], we present here a comprehensive study of the relationship between flies and yeast in wineries, using high-throughput, amplicon sequencing of the fungi associated with flies, coupled with well-established *Drosophila* behavior assays using both the yeast isolates and fly lines isolated from the same wineries. We demonstrate that *Drosophila* vector a distinct set of yeasts in wineries and exhibit a generally positive behavioral response towards commonly vectored yeasts. This suggests that the fly-yeast mutualism is not as specific as laboratory experiments indicate, and that flies interact with a diversity of yeast species in different ecological contexts.

## Results

### *Drosophila* vector wine yeasts in wineries

To identify the fungi vectored by flies, we collected adult *Drosophila* in three areas – fermentation tanks, cellars, and pomace piles – in four different wineries over two harvest seasons in the San Francisco Bay Area, California, USA (Figure 1A). In our initial harvest season, we collected in two wineries, one in Healdsburg, CA (HLD1) and the other in the Santa Cruz Mountains (SCM). We collected adult *Drosophila* every two weeks from May 2015 to November 2015. To expand our study in 2016, we collected flies in a four wineries, HLD1, SCM, HLD2 (also in Healdsburg, CA), and EBO (Orinda, CA) at a single time point from each winery in late September 2016 - early October 2016.

**Figure 1.**
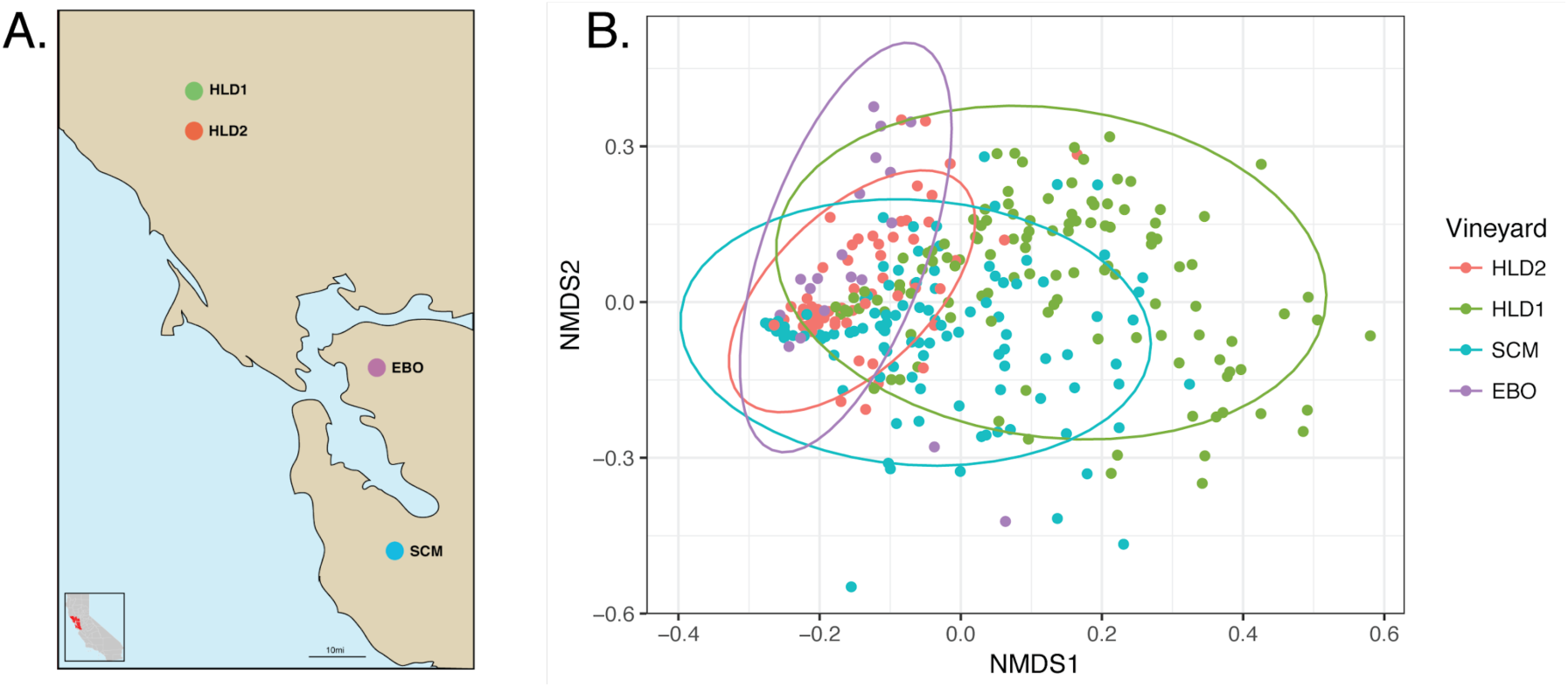
Fungal communities vectored by *Drosophila* are distinct between wineries. (A) Geographic locations of wineries sampled from 2015 and 2016 in the San Francisco Bay Area. (B) Bray-Curtis dissimilarity NMDS of fungal communities vectored by *Drosophila* in wineries (ADONIS: R^2^ = 0.129, *p*=0.001). Each sample was rarefied to 1000 sequences and is represented by a single point, color-coded by winery. Note, HLD2 and EBO have fewer samples because these wineries were only sampled in 2016.

DNA from whole, adult flies was extracted and short-amplicon sequencing targeting the universal fungal internal transcribed spacer region (ITS) was performed to characterize fungal community composition [31,32].

After quality filtering and processing, we clustered a total of 7,609,820 fungal ITS reads into 399 operational taxonomic units (OTUs) (Table S1). When rarefied to 1000 sequences per sample, the overall mean OTU richness per fly associated fungal community for each winery ranged from 19.741 +/- 7.673 to 39.771 +/- 10.537 OTUs (Figure S1, Table S2). We found that *Drosophila* species that were not *D. melanogaster* or its sister species *D. simulans* carried subtly but significantly different fungal communities (R_ANOSIM_=0.018, p<0.001, Figure S2) so we omitted these samples from our subsequent analysis. Removal of these samples resulted in 308 OTUs.

As expected, yeast species dominate the fungal communities vectored by *Drosophila*. The phylum Ascomycota, which includes many yeast species, represented the bulk of the fly-associated fungal taxa (average relative abundance: 95.6%). At the species level, fungal communities were dominated by *Hanseniaspora uvarum* (30.2% average relative abundance across all samples), *Pichia manshurica* (11.5%), *Issatchenkia orientalis* (10%), and members of the genus *Pichia* we could not identify at the species level (4.4%). Although *S. cerevisiae* is the dominant yeast in late stage fermentations, it was unevenly represented in the fungal communities vectored by *Drosophila*. Only 8.8% of the samples collected contained *S. cerevisiae* reads and of these samples, the relative abundance of *S. cerevisiae* ranged from 0.1% to 81.9% with no correlation to any particular winery or winery microhabitat.

In the laboratory, flies exhibit a strong attraction to fermentation volatiles, so it was unsurprising to find that drosophilds were associated with a range of fermentative yeast species commonly found in winery environments. However, the weak association with *S. cerevisiae* we observed was noteworthy given that *S. cerevisiae* is almost exclusively used in behavior assays investigating the fly-yeast mutualism in the laboratory. These results, corroborating Hoang et al, suggest that the use of *S. cerevisiae* in these assays may be irrelevant in a natural context [33]. Since we did observe a strong association between flies and non-*Saccharomyces* yeasts, we next asked if these associations were homogeneous across all winery *Drosophila* or dynamic across space and time.

### Nonrandom structure in the fungal communities associated with *Drosophila* between and within wineries

To elucidate the spatial and temporal patterns of the fly-fungi relationship, we asked if fungal community patterns could be distinguished between winery *Drosophila* populations. We found that the fungal communities vectored by flies are not randomly distributed and are significantly different between wineries (Figure 1B, Bray-Curtis R_ANOSIM_=0.129, p<0.001). These observations are consistent with what is known about the microbial communities present on wine grapes, which are predominantly defined by regional geography [34,35].

Unsurprisingly, fruit flies are found in vineyards and wineries are found predemoninantly in areas of active fermentation. To reflect this, we focused our drosophilid collections in three main areas in each winery: fermentation tanks, where primary fermentation occurs; cellars, where wine is aged; and the pomace pile, where grape berry waste is discarded. Adult flies are abundant at all of these winery microhabitats during wine production.

Within the HLD1 winery, we observed fungal community structure in the fungi vectored by flies between these three winery areas (Figure 2A, Bray-Curtis R_ANOSIM_=0.166, p<0.001). A single yeast species, *Hanseniaspora uvarum* (75.3%), dominated the fungal communities vectored by *Drosophila* collected from fermentation tanks. *Saccharomyces cerevisiae* (17.6%), *Hanseniaspora uvarum* (16.1%), *Pichia membranificiens* (14%), and *Penicillium brevicompactum* (10.7%) were overrepresented in the fungal communities carried by flies from the cellars while the pomace pile flies vectored primarily

**Figure 2.**
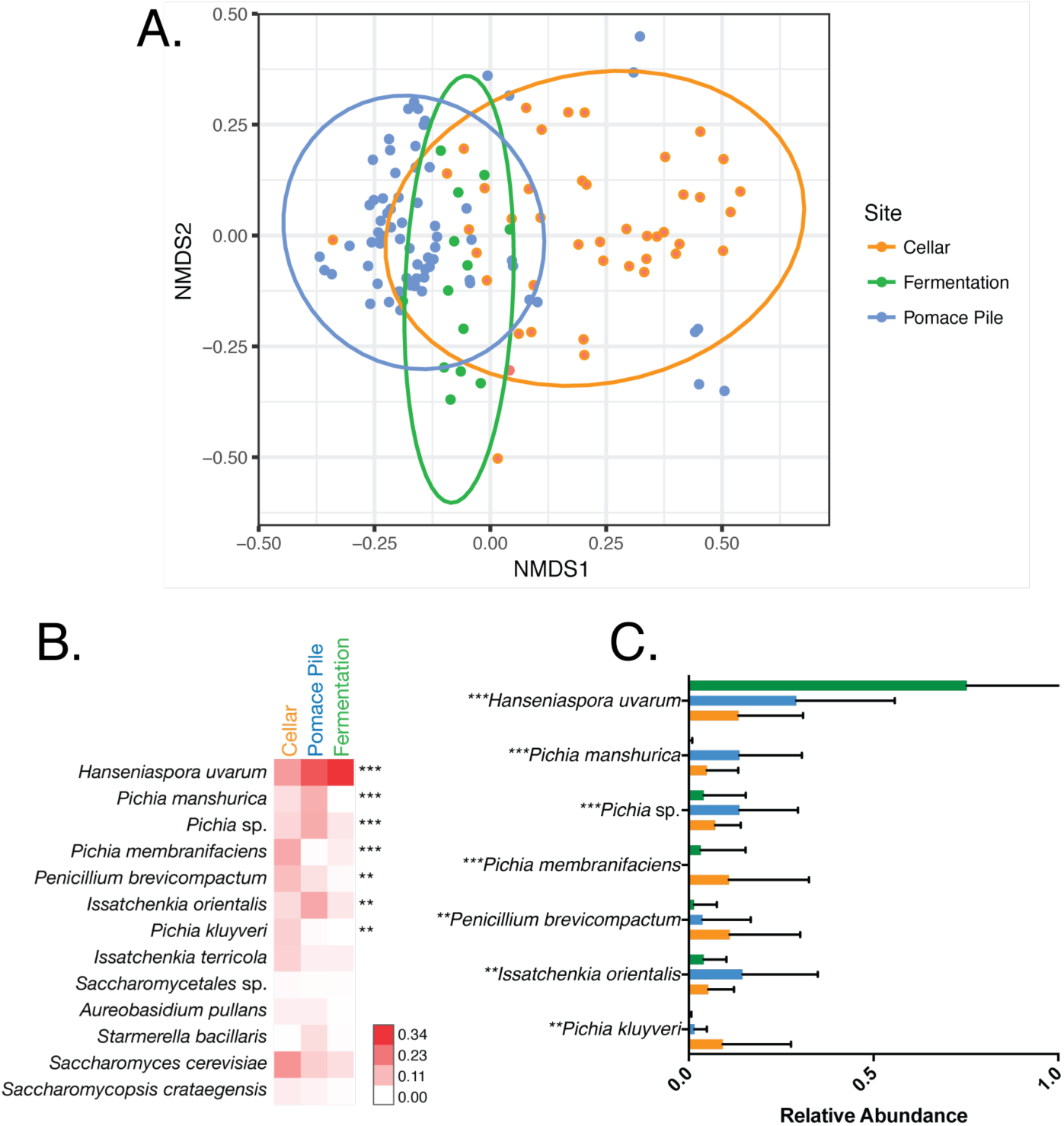
Within wineries, the fungal communities vectored by *Drosophila* are distinct between cellars, fermentation tanks, and pomace piles. The fly-associated fungal communities in winery areas are distinguished by the relative abundances of a few yeast species. (A) Bray-Curtis dissimilarity NMDS of fungal communities vectored by *Drosophila* in HLD1 in 2015 and 2016 (ADONIS: R^2^ = 0.166, *p*=0.001). Each sample was rarefied to 1000 sequences and is represented by a single point, color-coded by winery area. (B) Heatmap comparing the average relative abundances of all fungal species representing >1% of the total fungal community in each winery area. Each row represents a single fungal species. Stars to the right denote fungal taxa that have significantly different relative abundances between winery areas (one-way ANOVA with Bonferroni error correction, ns: not significant, *: p<0.05, **: p<0.01, ***: p<0.001, ****: p<0.0001.). (C) Bar graphs of the relative abundances of fungal taxa that are significantly different between winery areas.

### *Hanseniaspora uvarum* (28.2%), *Issachenkia orientalis* (14.6%), and *Pichia* species, such as *Pichia manshurica* (12.9%)

*Drosophila* in different winery areas carried many of the same fungal taxa but the relative abundances of these species distinguished fungal communities in one area from another (Figure 2B and 2C). Of these fungal species, the relative abundances of only six taxa were significantly different between the fungal communities vectored by flies in these specific winery areas: *Hanseniaspora uvarum, Pichia manshurica, Pichia membranificiens, Penicillium brevicompactum, Issatchenkia orientalis,* and *Pichia klyveri* (Figure 2C). All of these yeast species are commonly found in wineries and wineries [35]. Studies at other wineries have shown that winery equipment and processing surfaces harbor distinct microbial communities that change rapidly over time [34]. While we also observed distinct fly-associated fungal communities in different winery areas, the makeup of these communities do not fluctuate over time and do not completely mirror the previously identified fungal communities colonizing the fermentation tanks and cellar surfaces in other studies. Instead, *Drosophila* carry a subset of these taxa, suggesting that flies might play a role in shaping or maintaining the fungal community composition in these areas and only a subset of the yeasts from these fungal communities have a direct mutualistic relationship with flies.

### *Drosophila melanogaster* do not prefer yeast species representative of the winery area from which they were collected

We next asked how the fungal community structure between different winery areas is established. If flies actively modulate their associated fungal communities, we expected flies to prefer the yeast species characteristic of the fungal communities in the winery area from which they were established. We expected those preferences to manifest themselves in fly behaviors that are closely associated with the presence of yeast, such as olfactory attraction, oviposition, larval development, and longevity.

To test these hypotheses, we quantified the behaviors of four isofemale *Drosophila melanogaster* lines that were established from the three winery areas towards yeast isolates that were cultured and isolated from flies in the winery (Table 1). Founders of the fly and yeast lines were collected from the SCM and HLD1 wineries. We selected a panel of six yeast species that most strongly distinguished each winery area from the others (Figure 2C). Each winery area was represented by a single yeast species except for the pomace pile, which was represented by two because *Pichia manshurica* (isolate P2) was unable to ferment in liquid grape juice. We also included two controls in the yeast panel. *Issachenkia terricola* (yeast isolate CTLns) was included because it was vectored by all flies and did not distinguish one winery area from the others. Finally, *S. cerevisiae* (isolate CTLsc) was included as it is most often used in *Drosophila* behavior experiments in the laboratory and is generally attractive to *Drosophila melanogaster*, although we did not find that it was vectored frequently by wild flies in the winery.

**Table 1.** Fly lines and yeast isolates used in the behavior assays.

| Isolate/Fly Line | Type | Species | Vineyard | Site Collected | Year Collected | Month Collected |
| --- | --- | --- | --- | --- | --- | --- |
| CI | Yeast isolate | *Pichia klyveri* | HLD1 | Cellar | 2015 | August |
| FI | Yeast isolate | *Hanseniaspora uvarum* | SCM | Fermentation | 2014 | October |
| PI | Yeast isolate | *Issachenkia orientalis* | SCM | Pomace pile | 2015 | September |
| P2 | Yeast isolate | *Pichia manshurica* | HLD1 | Pomace pile | 2015 | August |
| CTLsc | Yeast isolate | *Saccharomyces cerevisiae* | SCM | Fermentation | 2014 | October |
| CTLns | Yeast isolate | *Issachenkia terricola* | SCM | Fermentation | 2015 | September |
| FermA | Isofemale fly line | *Drosophila melanogaster* | SCM | Fermentation | 2014 | October |
| FermB | Isofemale fly line | *Drosophila melanogaster* | SCM | Fermentation | 2014 | October |
| CellarA | Isofemale fly line | *Drosophila melanogaster* | HLD1 | Cellar | 2015 | August |
| PPA | Isofemale fly line | *Drosophila melanogaster* | HLD1 | Pomace pile | 2015 | August |

### Olfactory preference

Because *Drosophila* initially rely on olfactory cues to locate yeasts [16,36], we first tested differential attraction for the yeast species that distinguish the fly-associated fungal communities from different winery areas. We used a simple, olfactory-based assay previously developed in our lab [17] to quantify fly preference, and tested pairwise comparisons of a smaller yeast panel, with a single yeast representing each winery area.

Although we did find significant preferences between yeast species, these preferences did not reflect winery area and were variable between fly lines (Figure 3). In all lines except for FermA, *Pichia klyveri* (yeast isolate C1) was more attractive than both *Hanseniaspora uvarum* (yeast isolate F1) and *Issachenkia orientalis* (yeast isolate P1). Interestingly, FermA was the only fly line that was equally attracted to all three yeast species.

**Figure 3.**
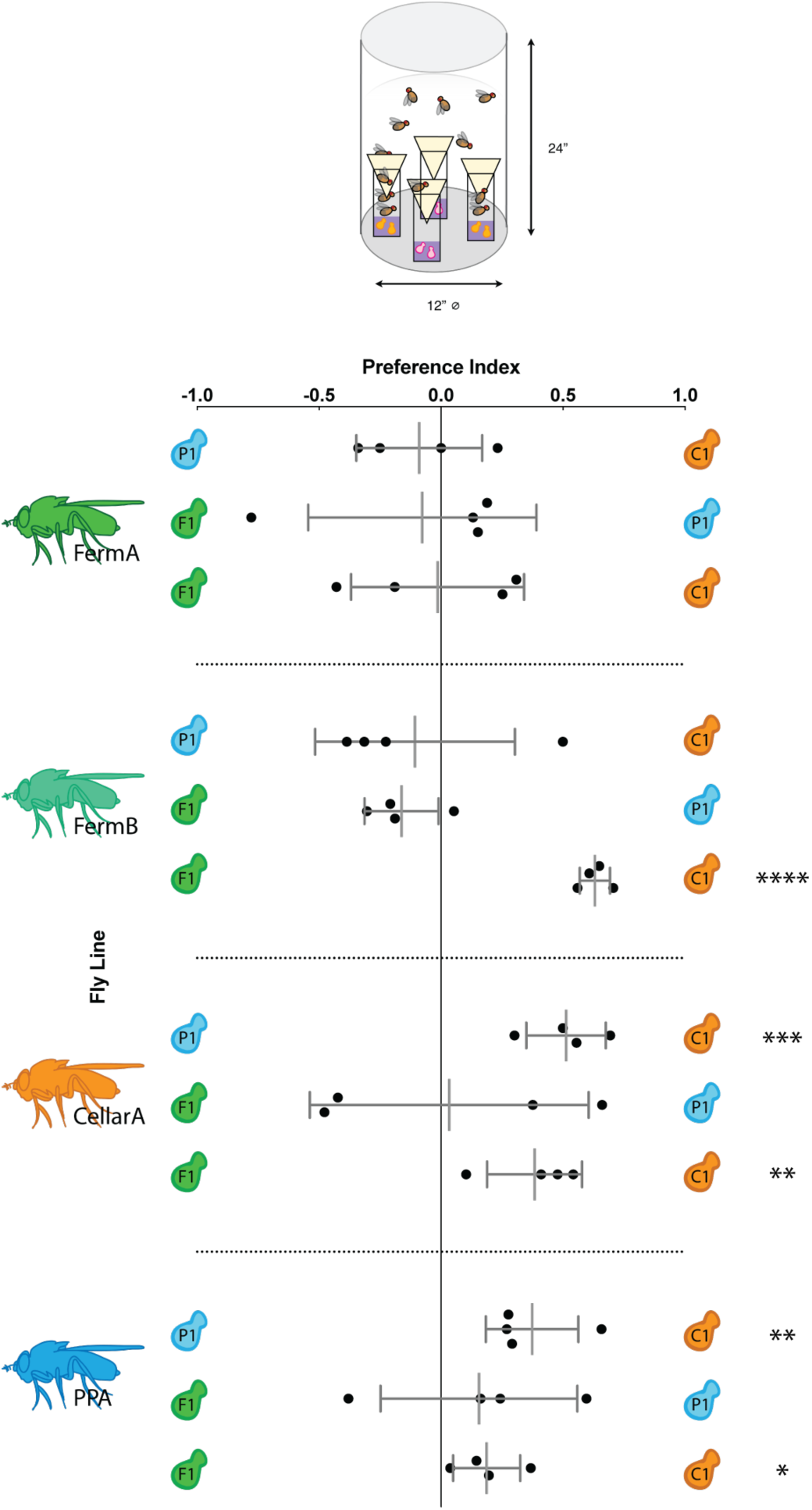
Based on olfactory cues, *Drosophila* do not prefer the yeast associated with their winery area. Preference was quantified using a trap-based choice assay in which adult flies were allowed to choose between traps with two different yeast strains. *Drosophila* lines tested are denoted by fly icons to the left and yeast species being compared are denoted by yeast symbols on the left and right axes. For a given comparison between yeast species A and B, preference index (PI) was calculated as: PI = (A-B)/(A+B), where A = total flies caught in A traps and B = total flies caught in B traps. A positive PI indicates a preference for yeast A, a negative PI indicates a preference for B, and a PI of 0 indicates no preference. Black dots indicate trial replicates. Short grey lines represent standard deviation and longer grey lines represent the mean of all trials. Stars to the left denote significantly different preferences between the two yeast species being tested (multiple t-tests with a Bonferroni correction, *: p<0.05, **: p<0.01, ***: p<0.001, ****: p<0.0001).

Fly lines did not prefer the yeast species that we had designated as representative of the fungal communities vectored by flies in their winery area. Instead, fly lines were more attracted to some yeasts over others with variability between lines. While *Drosophila* do exhibit some strong preferences between the yeast in the panel, these data suggest that *Drosophila* have a general attraction to yeast metabolites.

#### Oviposition

Where female flies choose to lay eggs is strongly coupled with offspring fitness [37,38]. We hypothesized that if *Drosophila* actively modulate the fungal communities they vector, female flies would prefer to oviposit on substrate inoculated with yeast representative of their winery area. To determine if this specificity exists, we tested female oviposition preference for sterile grape substrate or grape substrate inoculated with a single yeast species from our yeast panel (Figure 4).

**Figure 4.**
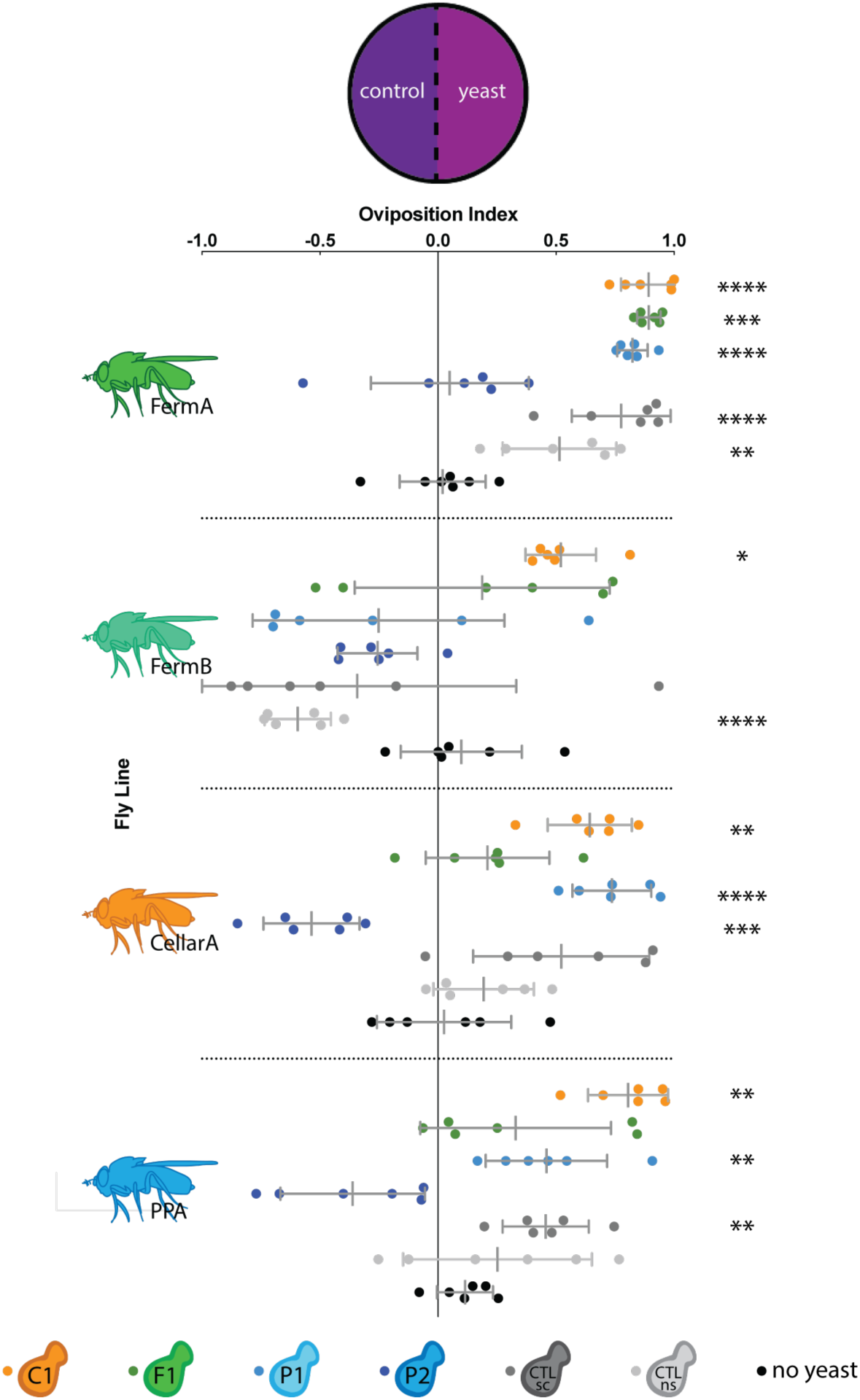
Most fly-associated yeasts elicit a generally positive oviposition response with variability between fly lines. Females from each fly line were given a choice between ovipositing on an agar plate with sterile grape juice or grape juice inoculated with a single yeast species. An oviposition preference index (OI) was calculated as: OI = (#eggs on yeast side - #eggs on control side)/(total #eggs deposited). A positive OI indicates an ovipositional preference for the yeast side, a negative OI indicates an ovipositional preference for the control side, and an OI of 0 indicates no preference. *Drosophila* lines tested are denoted by fly icons to the left. Individual replicates are represented by dots and are color-coded by yeast species. Short grey lines represent standard deviation and longer grey lines represent the mean of all trials. Stars to the left denote significantly different oviposition preferences between the two sides (multiple t-tests with a Bonferroni correction, *: p<0.05, **: p<0.01, ***: p<0.001, ****: p<0.0001).

*Pichia klyveri* (yeast isolate C1) was the only yeast species on the panel that elicited a significant, positive oviposition response (Figure 4) for all fly lines tested, which is consistent with olfactory preference results. In contrast, *Pichia manshurica* (yeast isolate P2) was the only yeast that elicited no preference or a negative oviposition response in all fly lines. Analysis of the metabolites produced by *Pichia manshurica* using gas-chromotography, mass-spectroscopy (GC-MS) showed that this yeast species is unable to ferment and produce volatile metabolites in liquid grape media (Figure S3). The lack of volatile attractants explains why *Pichia manshurica* elicits oviposition responses that mirror those of a sterile media control. Other yeasts in the panel, which fermented successfully, elicited intermediate ovipositional responses. These ovipositional preference patterns are not correlated with winery area. Instead, *Drosophila* seem to follow a general behavioral trend in response to the yeast panel, where some yeasts are more or less desirable oviposition substrates than others.

While all fly lines generally preferred to lay on yeast-inoculated media over sterile media (Figure 4), there was significant variation in oviposition response between fly lines (Table S3). For example, the two fly lines derived from the fermentation tanks, FermA and FermB, exhibited conflicting oviposition responses despite being collected at the same time. CellarA and PPA, which were collected in a different winery, winery area, and time, exhibited more similar, intermediate oviposition behavior. These data show that while there is natural heterogeneity in yeast volatile sensitivity between fly lines, the variation in ovipositional preference is not specific to winery area.

### Larval development

While oviposition substrate is important for larval fitness, larval development success and time is an indicator of nutritional health. *Drosophila* larva eclose both faster and more successfully when larval diet is supplemented with yeast [10,11]. To test if the yeasts associated with *Drosophila* in different areas of the winery have effects on larval development, we measured the development of winery fly lines when fed active monocultures of the yeast panel.

In corroboration with previous studies, we found that all fly lines develop more successfully on a diet supplemented with either live or dead yeast than on sterile media (Figure S4, Table S4). In fact, larvae grown on sterile media did not pupate at all, except for larvae from a single fly line, PPA, which only exhibited a 20% eclosion success on sterile media.

If a fly-yeast specificity existed in the winery, we hypothesized that larvae would develop faster and more successfully when supplemented with yeast species that are overrepresented in the fungal communities vectored by *Drosophila* in their winery area. For the subsequent statistical analyses, we omitted the no yeast and dead yeast controls from our dataset, as we were only interested in differential effects of the yeast species in our panel. We found that all yeast species on the yeast panel were equally suitable for *Drosophila melanogaster* development, with the exception of CellarA on *Pichia klyveri* (yeast isolate C1, Figure 5A). Only 64.4% of CellarA larvae supplemented with *Pichia klyveri* successfully eclosed compared to greater than 88% successful eclosion on all other yeast species in the panel (Table S4, p<0.0001 when compared to all other yeast species on panel). Interestingly, *Pichia klyveri* did not have a significantly different effect on eclosion success in other fly lines tested.

**Figure 5.**
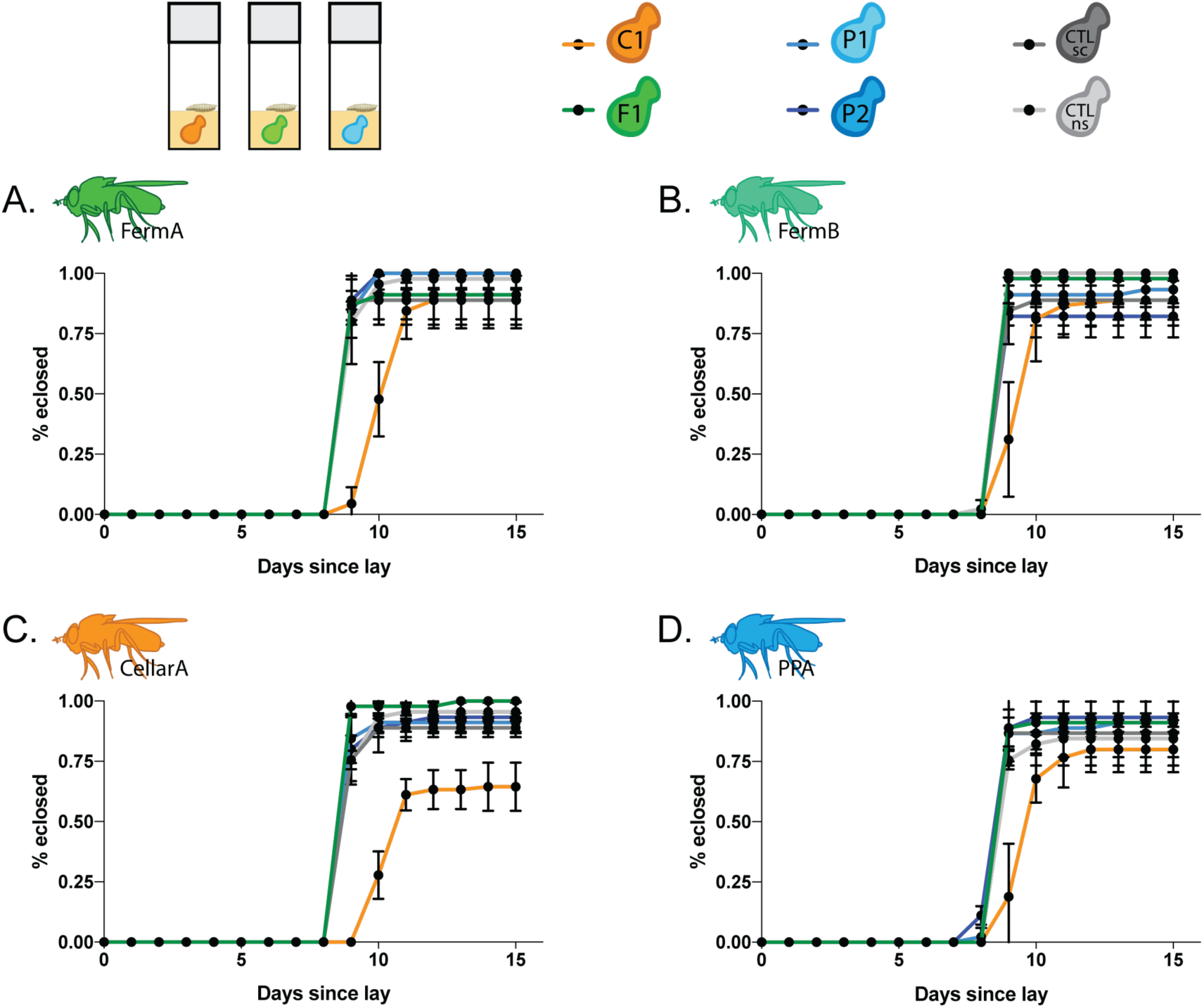
Some yeasts are more suitable for *Drosophila* development but do not follow a winery area specific pattern. Briefly, embryos from each fly line (A, FermA; B, FermB; C, CellarA; and D, PPA, as denoted by fly icons) were dechorionated with bleach to remove any maternally deposited microbes. Embryos were allowed to hatch on sterile media overnight and larvae were moved into vials containing a oneday, active monoculture of yeast. Eclosion time and success was monitored over time. Accompanying statistics for larval development time in Table S5.

Similar to the eclosion success, we found no significant differences in median development time between larva from all fly lines raised on active monocultures of the yeast species on our yeast panel except for *Pichia klyveri* (Figure 5, Table 2). While the median developmental time of all four fly lines on other yeast species was nine days, larva raised on *Pichia klyveri* had a delayed development time of 10-11 days (Table S5).

**Table 2.**
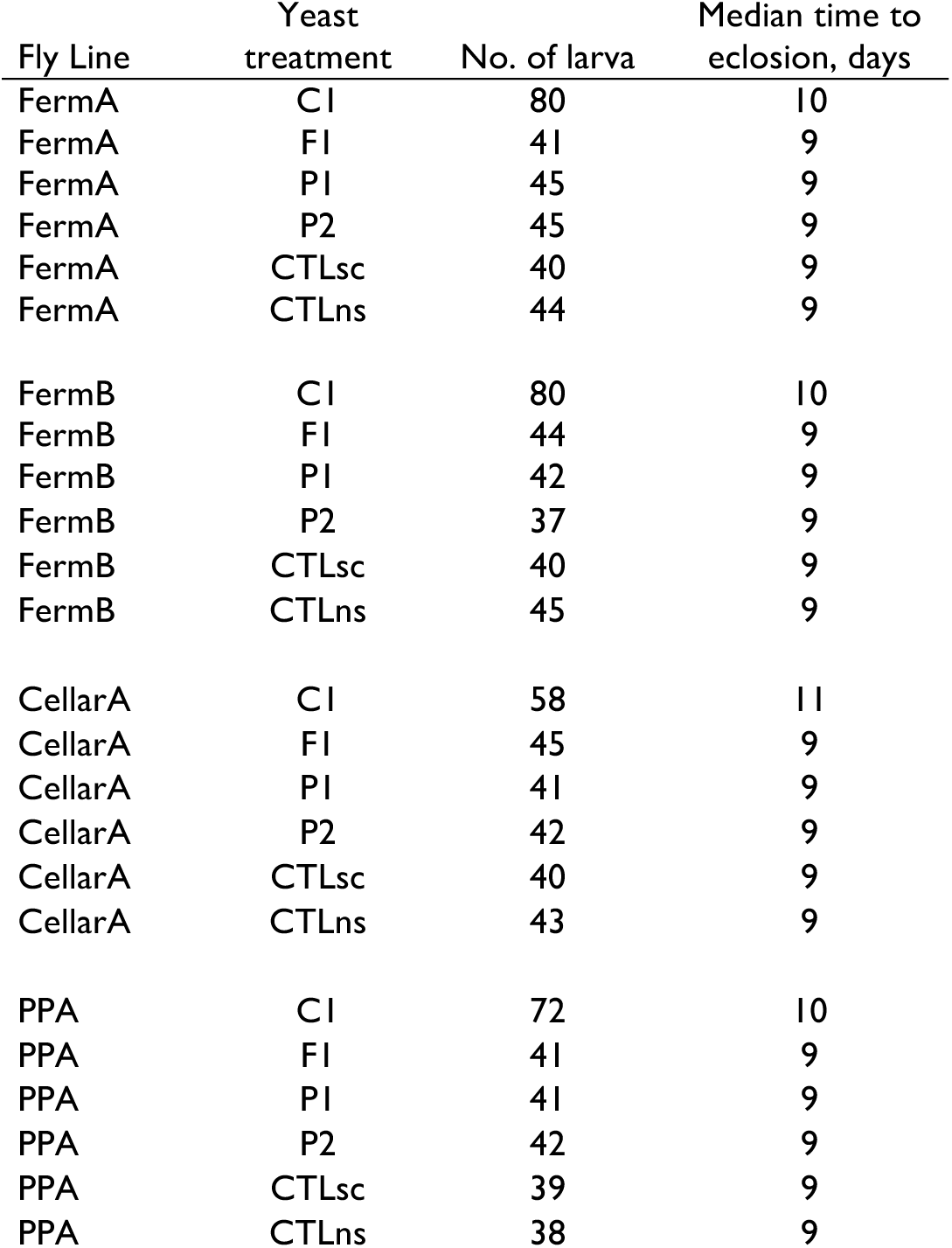
*Drosophila* larval development time from lay to eclosion when diet is supplemented with a single yeast species. Accompanying statistics in Table S5.

In our the behavior assays we described above, we showed that *Pichia klyveri* is both more attractive to adult flies and preferred as an oviposition substrate compared to other yeasts in the panel. So it is surprising that it has a negative effect on developmental timing. In addition, CellarA, a fly line established from a single female collected in a cellar, was the only line in which eclosion success was negatively affected by *Pichia klyveri*, which is commonly vectored by *Drosophila* in cellars. Together, these data suggest effects of yeast on *Drosophila* larval development are nonspecific to winery area. Instead, a broad range of yeasts are suitable for *Drosophila* larval development and while most are equally beneficial, some are less favorable than others.

### Longevity

Finally, we tested if the yeast species in our panel had winery area specific effects on *Drosophila* lifespan. Supplementing diet with yeast can rescue undernutrition and extend lifespan in *D. melanogaster* [39]. To measure if winery area yeasts have differential effects on *Drosophila* longevity, we maintained the adult flies that eclosed from the previous larval development assay on sterile media inoculated with the same yeast species throughout the lifetime of the fly. While these wild fly lines exhibit natural variation in lifespans (ANOVA, p<0.0001, Table S6), no particular yeast species had a significant effect on lifespan for a given fly line (Figure 6, Table 3). These results mirror those of the previously described olfactory preference, oviposition, and larval development assays.

**Table 3.** *Drosophila* lifespan when monoassociated with a single yeast species.

| Fly Line | Yeast treatment | No. of flies | Median lifespan, days |
| --- | --- | --- | --- |
| FermA | CI | 78 | 54 |
| FermA | FI | 35 | 49 |
| FermA | PI | 44 | 43 |
| FermA | P2 | 44 | 51 |
| FermA | CTLsc | 40 | 49 |
| FermA | CTLns | 40 | 50 |
| FermB | CI | 75 | 55 |
| FermB | FI | 41 | 61 |
| FermB | PI | 40 | 55 |
| FermB | P2 | 35 | 51 |
| FermB | CTLsc | 39 | 53 |
| FermB | CTLns | 45 | 54 |
| CellarA | CI | 57 | 58 |
| CellarA | FI | 42 | 53 |
| CellarA | PI | 36 | 58 |
| CellarA | P2 | 38 | 54 |
| CellarA | CTLsc | 35 | 48 |
| CellarA | CTLns | 38 | 56 |
| PPA | CI | 70 | 51 |
| PPA | FI | 33 | 45 |
| PPA | PI | 38 | 49 |
| PPA | P2 | 32 | 47 |
| PPA | CTLsc | 25 | 49 |
| PPA | CTLns | 36 | 51.5 |

**Figure 6.**
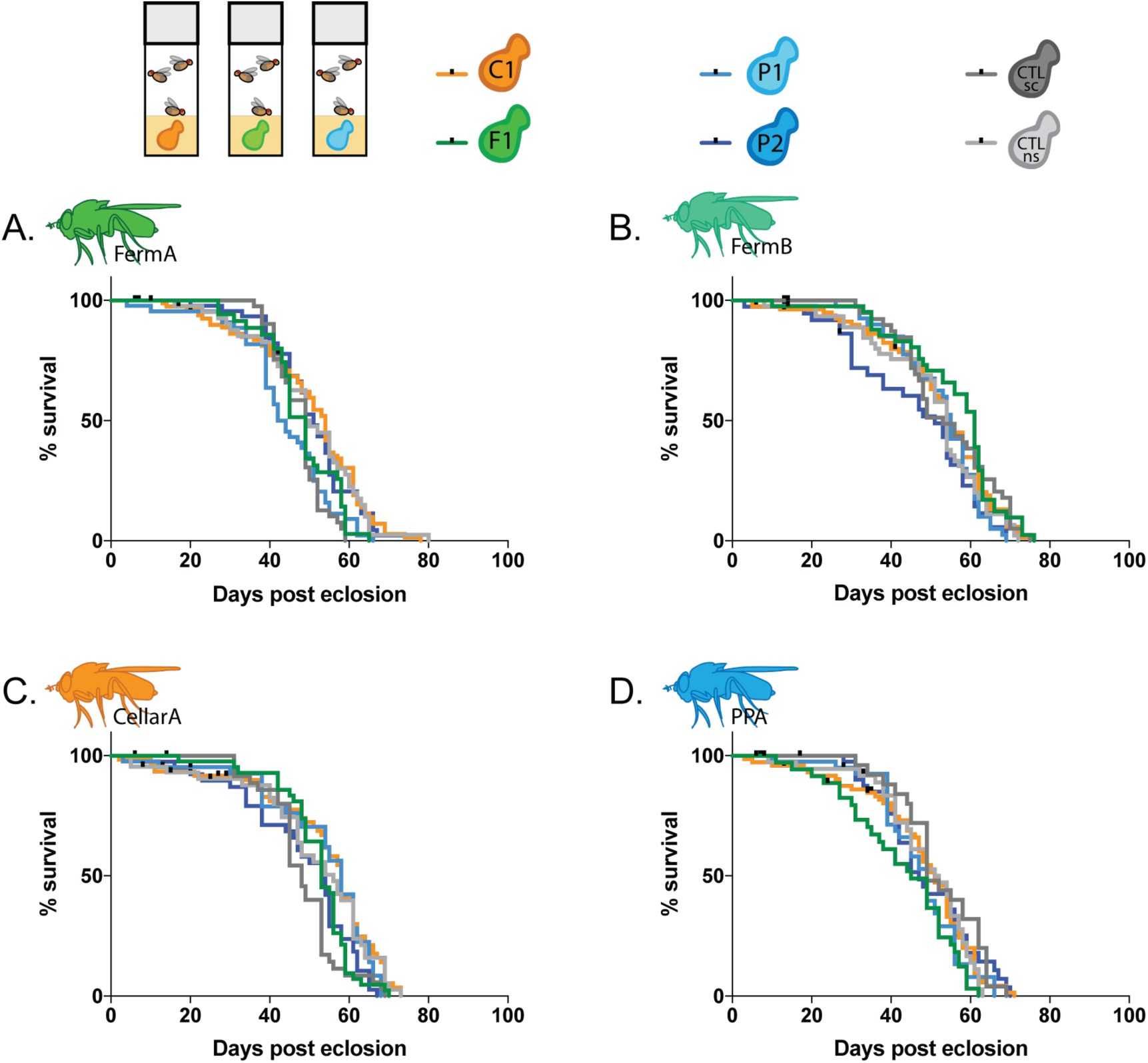
Fly-associated yeasts have no differing effects on *Drosophila* longevity. Survival curves of adult flies that have been associated with a single yeast throughout their entire lifecycle at 25C for each fly line (A, FermA; B, FermB; C, CellarA; and D, PPA, as denoted by fly icons). Adults that emerged from the previously described larval development assay were maintained in vials inoculated with a single yeast species and pushed onto new media twice a week. One day, active monocultures of yeast were used to reinnoculate vials once a week.

Overall, we found that while *Drosophila* exhibit variation in their behaviors towards some yeasts over others, these behaviors are not specific to winery area. These results consistently indicate that the structure in fungal communities vectored by *Drosophila* in different winery areas is not an obvious product of fly behavior.

## Discussion

Mutualisms are ubiquitous in nature, but understanding the factors mediating them requires insight into the natural context in which they operate and careful dissection of their parameters. The mutualism between fruit flies and yeast provide an ideal system in which to explore the constraints and specificity of a natural mutualism. In the laboratory, *Drosophila melanogaster* exhibit a remarkable ability to discriminate between different strains of *Saccharomyces cerevisiae* based solely on volatile profile [17,19] but whether these preferences are relevant in nature is less clear.

Our first goal in conducting this study was to determine if the fungal communities associated with *Drosophila* varied in a predictable way among the different fly-rich habitats in wineries. Our data clearly demonstrate that they do, with different abundances of a generally shared set of yeast species serving as a signature of fermentation tanks, cellars, and pomace piles.

This observation raises the more interesting question of how distinct fly-associated fungal communities are determined. There are, broadly, two possible, not mutually exclusive, explanations: flies could be passively sampling from environments with markedly different fungal populations, or flies could be selectively sampling fungi from these environments.

Using a set of well-defined *Drosophila* behavioral assays, we found that flies do not prefer the yeast species that distinguish the fungal communities in the winery areas from which they were sampled. Instead, the yeast species in our panel were generally attractive based on volatile cues, equally suitable for larval development, and have no differential effects on lifespan. Of course these laboratory experiments do not recapitulate the full range of behaviors manifested by flies in the wild, so we can not rule out the possibility that the stratification of fly-associated yeast communities with habitat is a direct result of fly behavior. However, our data suggest this is unlikely.

The alternative explanation is that flies are sampling passively from environments that are colonized by different fungal populations. The growth conditions experienced by yeast in different winery habitats likely influences yeast community structure. The substrates available - crushed must, spillage, or discarded grape fruits and stems - while related, vary in nutrient level enough to subtly favor the growth of some yeast species over others. Environmental factors, such as temperature, humidity, and sunlight, are likely to result in an even more pronounced effect on yeast species composition. Finally, the rate at which yeast species enter and exit each of these systems almost certainly varies.

Before harvest, the surface of grape berries are dominated by filamentous, non-fermentative fungi, presumably due to the inaccessibility of the sugar-rich flesh of fruits to fermentative yeast species [35]. However, once the grapes are harvested and crushed, their fungal communities are dominated by yeasts, reflecting the rapid growth of yeasts once their nutrient rich insides are exposed. Some of these yeasts were likely present on grape skins prior to harvest while others were probably deposited by flies.

Despite an abundance of sugar-rich substrates, well-maintained vineyards have relatively few *Drosophila*, likely reflecting the lack of fermentative yeast species before harvest. However, *Drosophila* become abundant as soon as grapes are harvested, and, even when measures are taken to control them, maintain healthy populations wherever crushed grape must, fermenting wine, and associated waste products are available.

Many observations, including this study, have shown that flies carry diverse fungal populations, and that they can transfer these populations to new substrates [13,25,27]. Moving forward it will be interesting to see how flies initialize and affect the dynamics of complex yeast populations, and to gauge the relative contributions of resident and vectored yeast in the process.

### Behavioral plasticity

The lack of behavioral specificity between flies and these yeasts indicates that a broad range of yeast species are beneficial to *Drosophila* and can serve adequately as a mutualistic partner. However, this does not imply that the fly-yeast mutualism is a weak interaction. Mutualisms occur on a spectrum of specific and nonspecific interactions, which require that each partner evolve traits that allow for more efficient interactions with the other partner. Development of these traits occurs on the continuum through evolutionary time.

In this study, we observed that while *Drosophila* do prefer specific yeast species, flies can vector and subsist on nearly all of them. This lack of specificity generally benefits all yeast species. However, we also observe that some yeasts are disproportionally vectored by flies and presumably, these yeasts are more preferred by flies. Yeast-produced fly attractants are mainly volatile, fruity esters hypothesized to serve an ecological function in the attraction of insects as vectors to new substrates [40]. This hypothesis is consistent with an independent studying showing that *S. cerevisiae* strains producing more attractive volatile cues achieve greater dispersal by *Drosophila simulans* [41] and indicates that ester production is under selection in yeast to ensure a close association with flies.

Conversely, *Drosophila* generally benefit from all yeast species, which is important in a naturally fluctuating environment where there are seasonal changes in food sources. The natural variation in yeast sensitivity and preference between wild fly lines we observed in this study likely reflects the known presence of dynamic, seasonal variants in natural *Drosophila* populations as a result of this seasonality [42,43]. As in other types of mutualisms, this relaxed specificity likely allows *Drosophila* to persist in a dynamic environment where food sources and temperatures shift throughout the year.

Most yeast species were adequate food sources for *Drosophila* larvae and adults, however, some yeast species are inherently more beneficial than others. It is advantageous for flies to be able to discriminate between and select yeast species that increase fitness. These adaptations could explain why we observed very specific olfactory and egg laying preferences between some naturally associated yeast species in this study and between strains of *S. cerevisiae* in the laboratory in previous studies.

However, flies’ ability to discriminate between yeast species is evolutionarily dynamic such that less beneficial yeasts can evolve traits that are more attractive to flies without any conferred benefit to fly fitness. The yeast species, *Pichia klyveri* from this study seems to exemplify this. *P. klyveri* was more attractive in the olfactory preference and oviposition assays than other yeasts but was the only yeast species that had a negative effect on larval fitness. Together, these results suggest that *P. klyveri* can produce volatiles that are more successful in attracting *Drosophila* than other yeast species, thereby increasing the chances of *P. klyveri* being vectored to new substrates, without any conferred benefit to the reproductive fitness of flies. This is likely a result of reciprocal selection between flies and yeast and the evolution of these traits, which allow partner species to interact with each other, is essential to the function of mutualisms.

While the fly-yeast mutualism can be specific, in a natural context, there are a diverse set of yeast species that attract *Drosophila* and influence the fly lifecycle positively. Extreme specialization is not a rule of mutualisms, in fact, most mutualisms tend to be nonspecific [44,45]. Therefore, the fly-yeast mutualism is a useful and promising one for continuing to define the parameters of mutualisms. This study leaves several questions about the constraints governing the fly-yeast relationship in nature. The yeast used in this study were commonly vectored by *Drosophila* so assaying the attractiveness of non-fly or rarely associated yeast species can reveal more about the specific yeast-produced metabolites influencing fly behavior. In the future, exploring the genetic basis driving yeast preferences in different *Drosophila* populations may elucidate more about the adaptations in play. Finally, wineries are habitats where both flies and yeast co-occur in large numbers and while this was advantageous for the goals of our study, we recognize that there are more natural contexts in which to study the fly-yeast mutualism. We hope that future studies will continue to study this relationship in less developed ecosystems.

## Materials and Methods

### Field collections of *Drosophila* and yeasts

All the wineries participating in this study practice organic farming and use spontaneous fermentations in winemaking. Adult *Drosophila* were collected in individual, sterile vials by direct aspiration or netting in cellars, fermentation tanks, and pomace piles. In 2015, flies were collected from two wineries in Healdsburg, California (HLD1) and the Santa Cruz Mountains, California (SMC) every two weeks from May 2015 – November 2015. During the 2016 harvest season, flies were collected from all four winery sites (Figure 1A) at a single time point from late September to early October. Flies were immediately transported back to the laboratory and processed within four hours of collection. Upon arrival at the laboratory, we documented sex and grouped samples into either *D. melanogaster/D. simulans* or other *Drosophila* species by eye.

### Isolation and identification of yeasts for behavior assays

Roughly one third of the total flies collected were cooled in individual vials on ice for two minutes to reduce activity and placed onto 5% YPD agar plates (Table S7). Roughly equal numbers of males and females were sampled. Flies were allowed to walk on plates overnight at ambient room temperature to deposit yeasts on plates and were aspirated off of plates in the morning. Yeast deposited on the plates were allowed to grow at ambient room temperature for 3-4 days. Single colonies, representing every yeast morphology present, were picked by eye and streaked onto fresh 5% YPD agar plates. If plates were overgrown with mold or single colonies were unable to be picked, a subsequent isolation was performed on a fresh plate. Isolated colonies were allowed to grow at ambient room temperature and stored at 4°C until identification by PCR. Original plates were kept for an additional three days after picking to ensure slower growing yeast were sampled.

Yeast colonies were identified by Sanger sequencing of the internal transcribed spacer region (ITS) [31]. Colony PCR reactions were performed in 25uL reaction volumes as follows: 12.5uL GoTaq Colorless Master Mix (Promega), 2uL of ITS1 and ITS4 primer at 10uM, 8.5uL nuclease-free water (Promega), and colony spike-in. Reaction conditions were as follows: 95°C for 5 min followed by 35 cycles of 95°C for 30 s, 53°C for 30 s, and 72°C for 60 s, and a final extension of 72 °C for 4 min (Jeremy Roop, personal communication). Amplification was verified on an agarose gel before being sent for Sanger sequencing (ELIM Biopharmaceuticals). Resulting sequences were trimmed for quality and then identified using BLAST (NCBI). Hits with an identity score greater than 98% were documented. After positive identification, yeast isolates were frozen as −80°C glycerol freezer stocks using standard protocol (Methods in Yeast Genetics, 2005) until use in behavior assays.

### Establishment of isofemale fly lines for behavior assays

Female *Drosophila* used for the yeast collection described above were aspirated onto fresh fly media (standard cornmeal-molasses-yeast medium, Bloomington Drosophila Stock Center) and used to establish isofemale lines. Each line underwent three rounds of intensive inbreeding where one to three virgin females were mated to three male siblings. After intensive inbreeding, lines were inbred at least 25 generations (10-40 female and male siblings) before being used in behavior assays.

### DNA extraction for amplicon study

The remaining flies collected (about 2/3 of flies) were immediately frozen and stored in individual, sterile 1.5mL microtubes for amplicon analysis. DNA was extracted following QIAGEN’s QIAamp Micro Kit tissue protocol with the modifications briefly described below [46]. After the overnight digestion with proteinase K, samples were bead beat (0.5 mm Zirconium beads, Ambion) in 200uL of WLB (Table S7). The samples were beat beat (MoBio) twice for one minute at 4C with a 30 second break in between, spun five minutes at ∼14,000xg, and the supernatant was transferred to a new tube. Beads were resuspended in 1ml of buffer WLB and beat again an additional minute, spun down, and supernatant was pooled. Finally, beads were washed once more with 1mL of buffer WLB, spun down, and supernatant was pooled. The pooled supernatant was spun down to pellet any beads and transferred to a clean microtube. One ug of carrier RNA (QIAGEN) dissolved in buffer AE was added to the supernatant before proceeding to ethanol precipitation and elution per the manufacturer’s protocol. DNA samples were quantified (Qubit dsDNA HS assay kit, ThermoFisher Scientific) and stored at −20C.

### Amplification and library construction

Fungal communities were characterized by amplifying the universal fungal internal transcribed spacer region I (ITSI) using BITS and B58S3 primers designed by Bokulich and Mills [32]. Each forward BITS primer includes a unique 8bp barcode connected to the universal forward primer with a 2bp linker sequence (sequences generously provided by Bokulich and Mills).

The following pre-PCR steps were carried out in a biosafety cabinet. The biosafety cabinet and laboratory supplies used were cleaned at the start of each day as follows to minimize PCR contamination: 10% bleach for 20mins, rinsed with autoclaved MilliQ water, 3% hydrogen peroxide for 10mins, and UV lamp for at least 5mins. PCR reactions were carried out in triplicate following the protocol previously used in [34] and described below.

For a single PCR reaction, reagents were added in the following order: 12.5uL GoTaq Colorless Master Mix (Promega), 2uL of B58S3 primer at 10uM, 5.5uL nuclease-free water (Promega), 2uL of BITS primer at 10uM, and finally 5–100 ng DNA template. Reaction conditions were as follows: 94 °C for 3 min, 35 cycles of 94 °C for 45 s, 50 °C for 60 s, and 72 °C for 90 s, and finally extension of 72 °C for 10 min [34]. PCR reactions were performed in 96 well plates with sample columns randomized between replicates to control for potential well biases. Both positive mock cultures (based on design described in [47], Table S8) and negative controls (from extraction and amplification steps) were randomized in the PCR plates among real samples.

PCR replicates were pooled and cleaned using AMPure XP magnetic beads according to manufacturer’s protocol (Beckman Coulter). Pooled PCR products were quantified using the Qubit dsDNA HS assay kit and the samples on a single PCR plate were pooled at 30ng equimolar concentration. Any samples with concentrations of <1ng/uL were omitted. Negative control samples with concentrations of <1ng/uL were pooled to the largest volume of real samples. After pooling, each pool was cleaned and concentrated according to the manufacturer’s protocol (Zymo Clean and Concentrator), eluted in 22uL of sterile water, and quantified with the Qubit dsDNA HS assay kit.

Illumina sequencing libraries were prepared for each PCR plate pool using TruSeq RNA v2 kit (Illumina) beginning at the A-Tailing step of the manufacturer’s protocol using at least 200ng of starting material. A different Illumina index was used for each PCR plate pool. Libraries were verified and quantified using the Qubit dsDNA HS assay kit and the Agilent 2100 Bioanalyzer (Agilent Technologies). Up to four libraries were pooled and sequenced on a single 250bp paired-end Illumina MiSeq lane. Samples from 2015 were sequenced at the Vincent J. Coates Genomics Sequencing Laboratory (UC Berkeley) and the 2016 samples were sequenced at the UC Davis DNA Technologies Core. To control for the change in sequencing services, we resequenced a library from 2015 at UC Davis with 2016 samples and achieved very replicable results (Table S8, Figure S6).

### Data analysis of amplicon study

We followed many of the same processing steps outlined in [35,48] as we used the primers designed in these studies and expected the fungal communities of *Drosophila* in wineries to be similar to the fungal communities in wineries and breweries. Raw and quality filtered sequence counts from the following steps are summarized in Table S1. Raw read pairs were merged using BBMerge (https://jgi.doe.gov/data-and-tools/bbtools/), demultiplexed in QIIME v1.9.1 [49] and primer sequences were trimmed using cutadapt [50]. Resulting reads were quality filtered in QIIME as follows: any read less than 80bp was removed, any read with more than 3 consecutive bases with a quality score <19 was removed, and any chimeric sequences were filtered against the UCHIME chimera reference dataset v7.1 [51]using the union method.

Open reference OTU picking was performed in QIIME using the UCLUST method [52]against a modified UNITE database [53,54]with a threshold of 97% pairwise identity. Sequence alignment and treebuilding were suppressed and taxonomy was assigned using BLAST (NCBI). After OTU picking, positive control mock culture samples and OTUs with ‘no blast hit’ were filtered. Using R version 3.2.4 [55], negative controls were removed, max sequence counts of all negative control OTUs were calculated, and then subtracted from all real samples to account for spurious sequences produced from possible PCR, sequencing, or spillover contamination [47,56]. Finally, an OTU threshold of 0.001% was applied [35,48].

Alpha diversity measurements of observed OTU richness were calculated and visualized in QIIME to reveal that all samples had been sequenced to saturation (Figure S1). Community analyses were conducted using the vegan [57] and biom [58] packages in R. To determine relationship between the fungal communities vectored by *Drosophila*, samples were evenly subsampled to 1000 reads per sample and Bray-Curtis dissimilarity was calculated and visualized with nonmetric multidimensional scaling (NMDS) using ggplot2 [59]. ADONIS was used to calculate the relative effects of factors that distinguished fly-associated communities from others. The fungal communities associated with *Drosophila simulans*/*Drosophila melanogaster* samples were significantly different than those of other *Drosophila* species (Figure S2) so other *Drosophila* samples were filtered out.

The relative abundances of fungal taxa in different winery areas were clustered using hierarchical clustering by taxa (Cluster 3.0) and visualized in Java TreeView and Prism 7 (GraphPad). To identify fungal taxa with significantly different relative abundances between winery areas, OTUs were collapsed by species name and the Kruskal-Wallis test was employed (with Bonferroni correction) in QIIME.

### Olfactory preference assay

A custom trap-based olfactory preference assay previously designed by Schiabor et al 2014 was used to measure *Drosophila* olfactory preference [17]. Yeast species were plated onto agar grape juice plates (Table S7) and grown at 30C for 22 hours. The following day, plates were removed from the incubator, fitted with a custom printed lid, and secured with Parafilm. Lids were topped with a 50mL conical centrifuge tube (Falcon) with the end removed and covered in mesh. A funnel was fashioned from 150mm filter paper (Whatman, 150mm, Grade 1) with a 5mm hole snipped off the tip and secured to the top of the centrifuge tube with tape.

*Drosophila melanogaster* lines were raised at room temperature (21-23C) on standard cornmeal-molasses-yeast media and aged at room temperature for at least four days under ambient lighting conditions (i.e. adjacent to a window) before being used in behavior assays. One hundred and twenty 4-10 day old mixtures of male and female flies were anesthetized with CO2 and allowed to recover on cornmeal-molasses-yeast media for 2 hours before being used in behavior assays.

Pairwise comparisons of yeast were used to assay for olfactory preference. Two traps for each yeast species were placed into behavior arenas (*Drosophila* population cages, 24” x 12” clear acrylic cylinders, TAP plastics) and fitted with netting (Genesse Scientific) as shown in Figure 3 and Figure S5A. All four possible orientations of plates within the arena were tested to control for potentially confounding environmental variables such a light (Figure S5B).

Flies were introduced into behavior arenas at 3pm and allowed survey traps. After 18 hours, traps were removed from the arena and the number of flies in each trap were counted, sexed, and recorded. Flies were only used in behavior assays once and were discarded after counting. A preference index was calculated from the number of flies in each trap as follows:

For **A** = total number of flies in traps baited with Yeast A

For **B** = total number of flies in traps baited with Yeast B

Preference Index = (**A** - **B**)/(**A** + **B**)

A positive PI indicates a preference for yeast A, a negative PI indicates a preference for B, and a PI of 0 indicates no preference. Multiple t-tests with Bonferroni correction were executed in Prism 7 and used determine which yeast preferences were significant.

### Oviposition assay

The egg laying assay in this study was adapted from Joseph et al 2009 [60] and Fischer et al 2017 [61]. At 10am, 75mL cultures of the yeast strain of interest in liquid grape juice (Table S7) were inoculated with 1.5mL of yeast cells suspended in sterile 1X PBS (Mediatech Inc) and diluted to OD_600_=1. These cultures, and a negative control, were grown shaking at 30C for 72 hours.

Oviposition assay cages were fashioned from polypropylene *Drosophila* bottles (6oz, square, Genesse Scientific) with the bottom cut out and covered with mesh. During acclimation, cages were capped petri dishes (35×10mm, Falcon) filled with grape agar premix (Genesse Scientific) and topped with yeast paste (Red Star).

Similar to olfactory preference assays, *Drosophila melanogaster* lines were raised and aged for four to ten days at room temperature on standard cornmeal-molasses-yeast media. Twenty four hours before the assay began, twenty non-virgin females were acclimated to oviposition cages at 25C. Cages were kept on the top shelf of a 25C incubator on a 12 hour light cycle, placed on the side, and positioned so the mesh bottom faced the back of the incubator and the plate faced the door of the incubator (Figure S5C).

At least one hour before the start of the behavior assay, acclimated cages were cleared by replacing the petri dish with a new grape agar premix plate without yeast paste and returned to 25C. After 72 hours of fermentation, cultures were removed from 30C, mixed 1:1 with a boiled water-agar solution (BD), and cooled to 65C to achieve a final agarose concentration of 1.6%. The lids of petri dishes (35×10mm, Falcon) were divided in half using laminated paper. Plates containing half uninoculated grape juice and half inoculated grape juice were created by pouring both sides simultaneously (Figure S5D).

Plates were cooled for 15 minutes at room temperature and the laminated paper was removed. Plates were immediately used in behavior assays by replacing the grape agar premix plate cage topper. *Drosophila* females were allowed to oviposit on plates for three hours before plates were removed for counting (usually from around 12noon to 3pm). Flies were only used once and discarded at the end of the assay.

An oviposition index (OI) was calculated from the number of embryos deposited on each side of the plate as follows:

For **Y** = total number of eggs oviposited on inoculated side

For **N** = total number of eggs oviposited on uninoculated side

Oviposition Index = (**Y** – **N**)/(**Y** + **N**)

A positive OI indicates an oviposition preference for the yeast side, a negative OI indicates a preference for the control side, and a OI of 0 indicates no preference for either side. Multiple t-tests with Bonferroni correction were used to determine whether the yeast tested elicited a significantly different oviposition preference in relation to the control. One way ANOVA was used to test for any differences in ovipostion responses between fly lines for a given yeast species. If ANOVA results were statistically significant, Tukey’s multiple comparisons test was used to identify the fly lines exhibiting ovipostion responses that were significantly different than other lines. These statistical analyses and those described in the methods following were implemented in Prism 7 with a significance cutoff of p>0.05.

### Gas chromatography – mass spectroscopy (GC-MS)

A subset of the oviposition plates used in the oviposition assays were also sampled by GC-MS in parallel with the behavior assays using a stirbar sorptive extraction (SBSE) and thermal desorption approach. Oviposition plates were placed in sterile 60 × 15mm petri dishes for headspace sampling. As previously described in [17], a conditioned, Twister stir bar (10 mm in length, 0.5mm film thickness, 24uL polydimethylsiloxane, Gerstel Inc) was suspended from the lid of the larger petri dish with rare earth magnets for 40 minutes at room temperature. The Twister bar was then dried using a Kimwipe, placed in a thermal desorption sample tube, topped with a transport adapter, and loaded onto sampling tray (Gerstel Inc).

Automated sampling and analysis was performed using the Gerstel MPS system and MAESTRO integrated into Chemstation software. Sample analysis was performed on an Agilent Technologies 7890A/5975C GC-MS equipped with a HP-5MS (30m × 0.25mm, i.d., 0.25micrometers film thickness, Agilent Technologies) column. Samples were thermally desorbed using the Gerstel Thermal Desorption Unit (TDU) in splitless mode, ramping from 30C to 250 °C at a rate of 120 °C/min, and held at the final temperature for 5 minutes. The Gerstel Cooled Injection System (CIS-4) was cooled to - 100C with liquid nitrogen before ramping to 250 °C at a rate of 12 °C/min and held for 3 mins for injection into the column. The injector inlet was operated in the Solvent Vent mode, with a vent pressure of 9.1473 psi, a vent flow of 30mL/min, and a purge flow of 6mL/min.

The GC oven temperature program was set to 40 °C for 2 min, raised to 140 °C at 4 °C/min, and finally raised to 195 °C at 15 °C/min and held for 10 min. A constant helium flowrate of 1.2 mL/min was used as carrier gas. The MSD transfer line temperature was set at 280C. The MS was operated in EI mode with the electron voltage set at autotune values. The detector was set to scan from 30 to 300amu at a threshold of 150 at a scanning rate of 2.69 scans/second. The ion source and quadrupole temperatures were set at 230C and 150C, respectively.

GC-MS data files were visually inspected using Chemstation and peaks were identified using the NIST O8 database. Datafiles were transferred, parsed, and analyzed using custom written Matlab scripts in [17]. Every chromatogram trace represents, at minimum, the average of 6 replicates.

### Larval development assay

In order to test the effects of each yeast species on larval development time and success, *Drosophila* larvae were raised on sterile, yeast-free media supplemented with a single yeast species. As in the previously described assays, *Drosophila melanogaster* lines were raised and aged for four to ten days at room temperature on standard cornmeal-molasses-yeast media. Twenty-four hours before embryo collection, at least 50 adults flies were acclimated to the oviposition assay cages capped with grape agar premix plates and yeast at 25C as described above.

At 9am the following morning, plates in oviposition cages were replaced with new grape agar premix with yeast plates to clear any hoarded eggs. After 30 minutes, clearing plates were replaced with new grape agar premix with yeast plates and flies were allowed to lay for two hours at 25C for embryo collection. After two hours, collection plate was removed and aged at 25C for two hours. Aged embryos were washed off plates with MilliQ water into a embryo wash basket fashioned out of a 50mL conical (Falcon) with the end cut off and the top of the lid removed and covered with thin mesh. In the wash basket, embryos were dechorionated with 30% bleach for three minutes, consequently removing any previously associated yeast. Embryos were washed with sterile, autoclaved MilliQ and then with sterile PBS-t (1X PBS, 0.5% triton). Using a sterile paintbrush, embryos were moved onto sterile agar plates and allowed to hatch at 25C overnight. For data analyses, this day was considered Day 0 of the assay.

On the same day at 9:30am, 5mL starter cultures of liquid 5% YPD (Table S7) were inoculated with the yeast species of interest and grown shaking at 30C. At 3:30pm, cultures were removed and diluted to OD_600_=0.5. *Drosophila* vials with sterile, yeast-free GB media (Table S7) were spotted with 50uL of diluted culture and grown at 30C overnight. Three replicate vials for each yeast species and each fly line were set up (Figure S5E).

At 11am the next day, embryo plates were removed from the incubator. Using a sterile paintbrush (dipped in 50% bleach, 75% ethanol, autoclaved MilliQ, and sterile PBS-t between each vial), 15 larvae were moved into each vial and allowed to develop at 25C. Due to the extensive setup and time constrains of fly development, we tested the larval development of all four fly lines for each yeast species on the panel in three groups over 2 months. In each group, a positive control on standard cornmeal-molasses-yeast media (dead yeast) and a negative control on sterile GB media with no yeast supplement was run in parallel with experimental conditions.

We opted to start our assays with larvae instead of embryos in an effort to control for any death after dechorionation. For data analyses, this day was considered Day 1 of the assay. Each day, vials were checked for emerged adults and randomized within the incubator until the assay was terminated on Day 16. Adults that eclosed successfully were moved into sterile GB media vials daily and subsequently used for the longevity assays described below.

Eclosion curves and statistics were plotted in Prism 7. To test whether larvae given any yeast eclosed more successfully than those given no yeast and whether some yeast species resulted in greater eclosion success than others, one way ANOVA followed by Tukey’s multiple comparisons test was used to calculate significance values for each yeast species and controls within each fly line.

### Longevity assay

To study the effects of each yeast species on the lifespan of *Drosophila*, adults that eclosed successfully from the larval development assay were fed a diet supplemented with the same yeast species throughout their lifetime. At 9am on Day 8 of the larval development assay, 5mL starter cultures of liquid 5% YPD were inoculated with the yeast species of interest and grown shaking at 30C. At 3:30pm, cultures were removed and diluted to OD_600_=0.5. *Drosophila* vials with sterile, yeast-free GB media (Table S7) were spotted with 50uL of diluted culture and grown at 30C overnight (Figure S5E).

As flies hatched off of the larval development assay, they were moved onto the inoculated media and checked every day. Each day, vials were randomized within the incubator to control for positional effects. Flies were maintained at 25C and pushed onto fresh media twice a week, once into sterile GB media and once into inoculated media prepared as described above.

Survival curves and statistics were plotted in Prism 7. One way ANOVA followed by Tukey’s multiple comparisons test was used to test whether different fly lines had significantly different lifespans. The effect of single yeast species on the lifespan of a single fly line was tested using one way ANOVA but found no significant differences.

## Funding and Acknowledgements

This work was funded through the National Science Foundation Graduate Research Fellowship Program and Howard Hughes Medical Institute. We are grateful for field assistance from Patrick O’Grady, Kelly Schiabor, Kyle Barrett, Elizabeth Roeske, Carolyn Elya, Addie Norgaard, Ciera Martinez, and Jesse Rau. We are also indebted to Nick Bokulich, David Mills, Angus Chandler, Rachel Adams, Sydney Glassman, Shana McDevitt, and Dylan Smith for their advice in fungal amplicon experimental design and data analysis. David Hembry and Will Ludington provided valuable feedback on the drafts of this manuscript. None of this would have been possible without the generosity of Ridge Vineyards, DaVero Farms & Winery, Les Lunes Wine, and Populis Wine with special thanks to Shun Ishikubo, Eric Baugher, Mike Bairdsmith, and Will Thomas.

## Supplementary Figures and Tables

**Figure S1.**
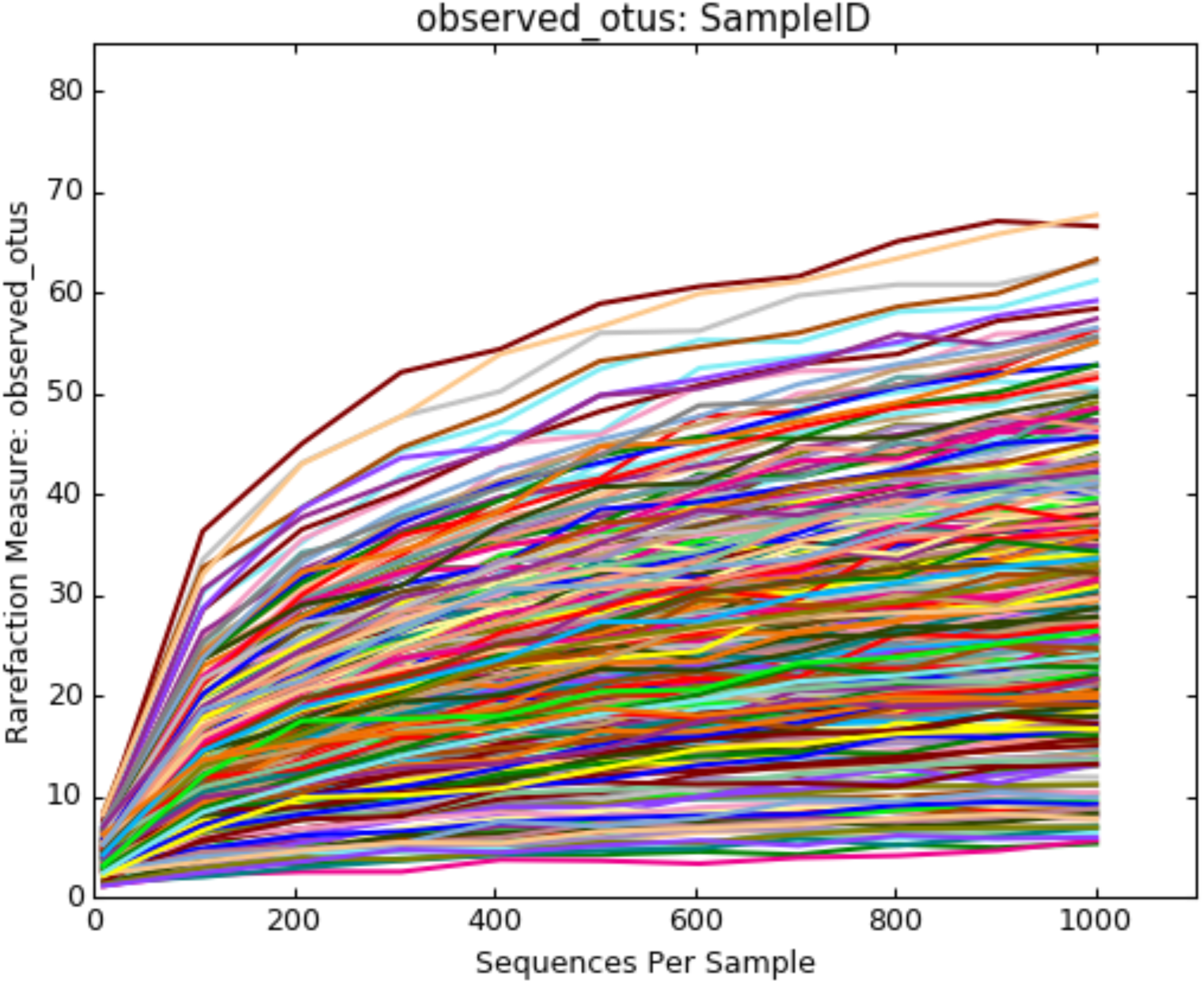
Alpha diversity richness QIIME rarefaction curves.

**Figure S2.**
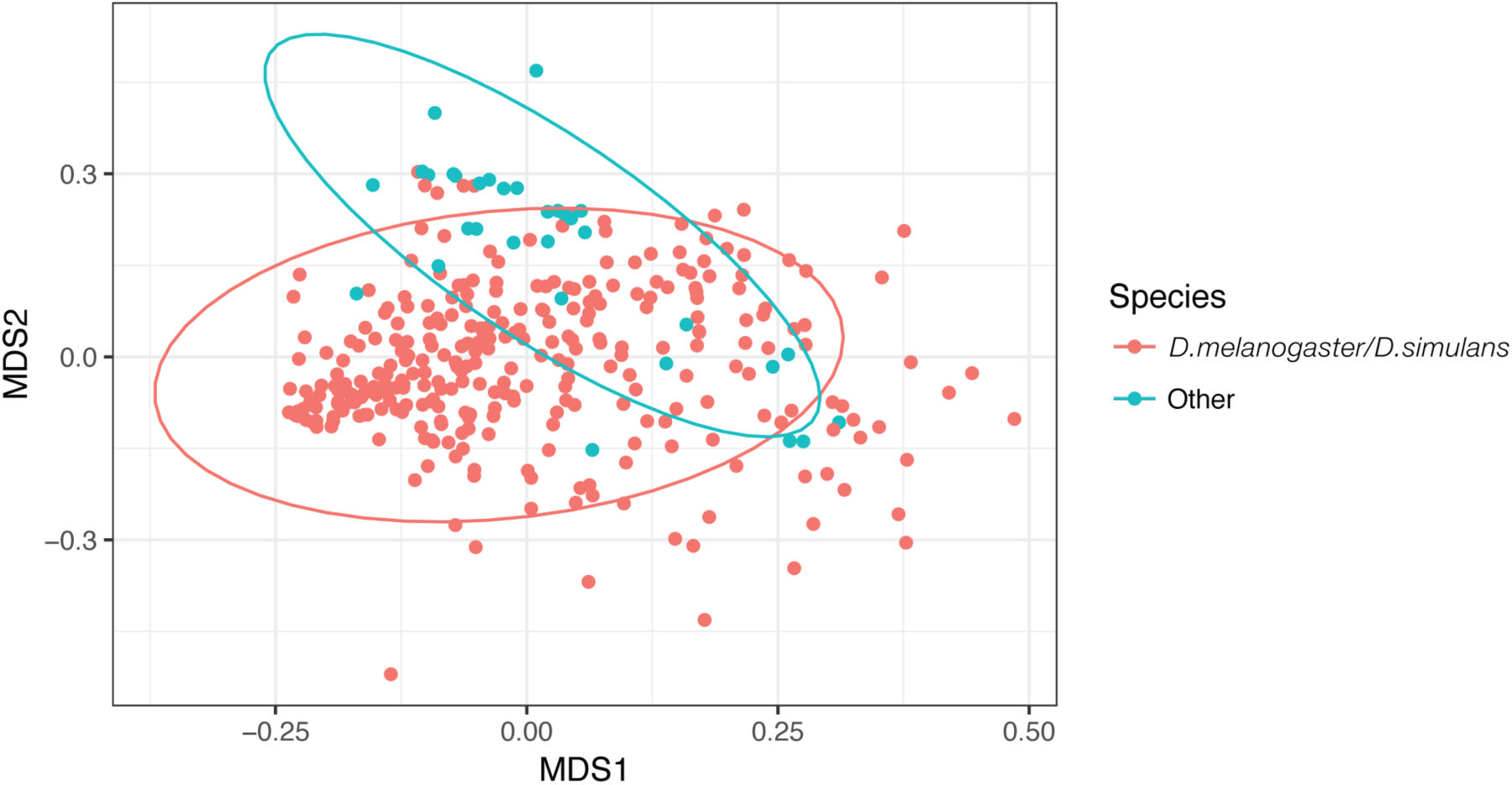
The fungal communities vectored by *Drosophila melanogaster/Drosophila simulans* are different than other *Drosophila* species (ADONIS: R^2^ = 0.018, *p*=0.001). Bray-Curtis dissimilarity NMDS of fungal communities vectored by *Drosophila* in all vineyards in 2015 and 2016. Each sample was rarefied to 1000 sequences and is represented by a single point, color-coded by species.

**Figure S3.**
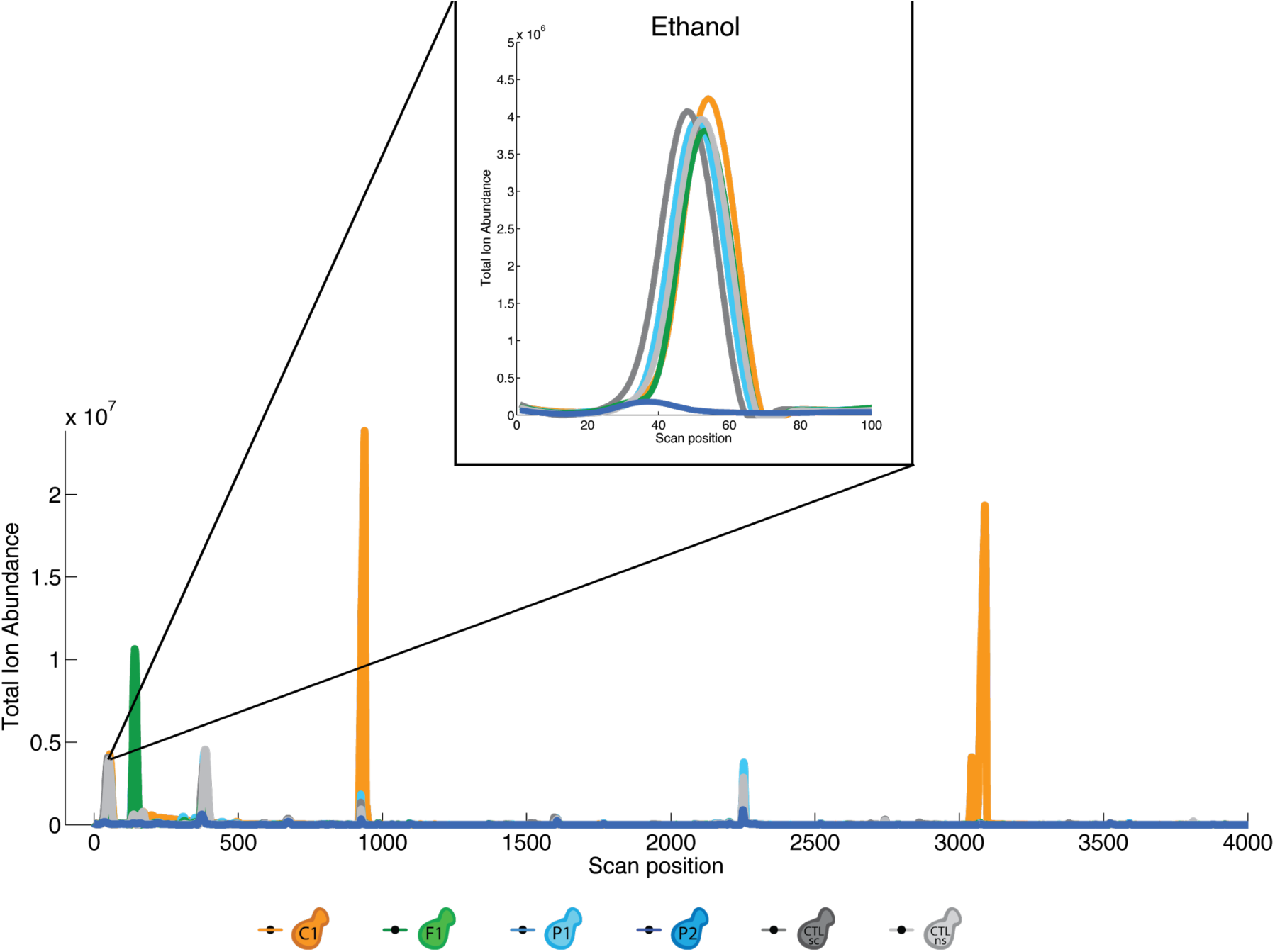
*Pichia manshurica* (yeast isolate P2) does not ferment well in liquid grape juice and when measured by GC-MS, produces very low levels of ethanol (inset) and other volatile metabolites compared to other yeast species on the panel. Each line represents the average of eight GC-MS replicates for a given yeast species. Replicates were sampled for GC-MS in parallel with oviposition assays.

**Figure S4.**
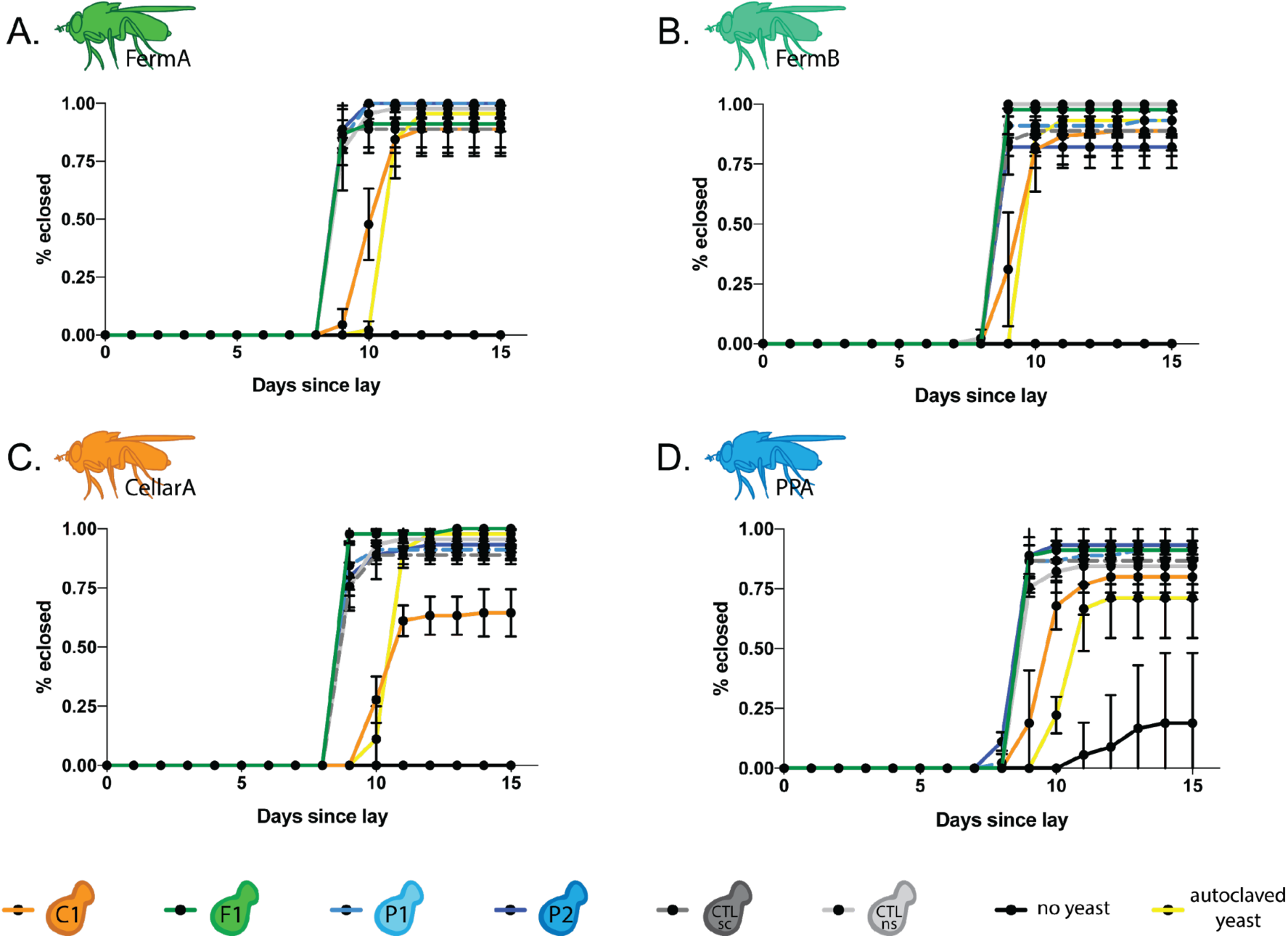
*Drosophila* larvae supplemented with any species yeast species, dead or alive, develop more successfully than those fed no yeast at all. Note that data for larvae that were not given yeast only exist for PPA line because no larvae eclosed without the addition of yeast in any other lines.

**Figure S5.**
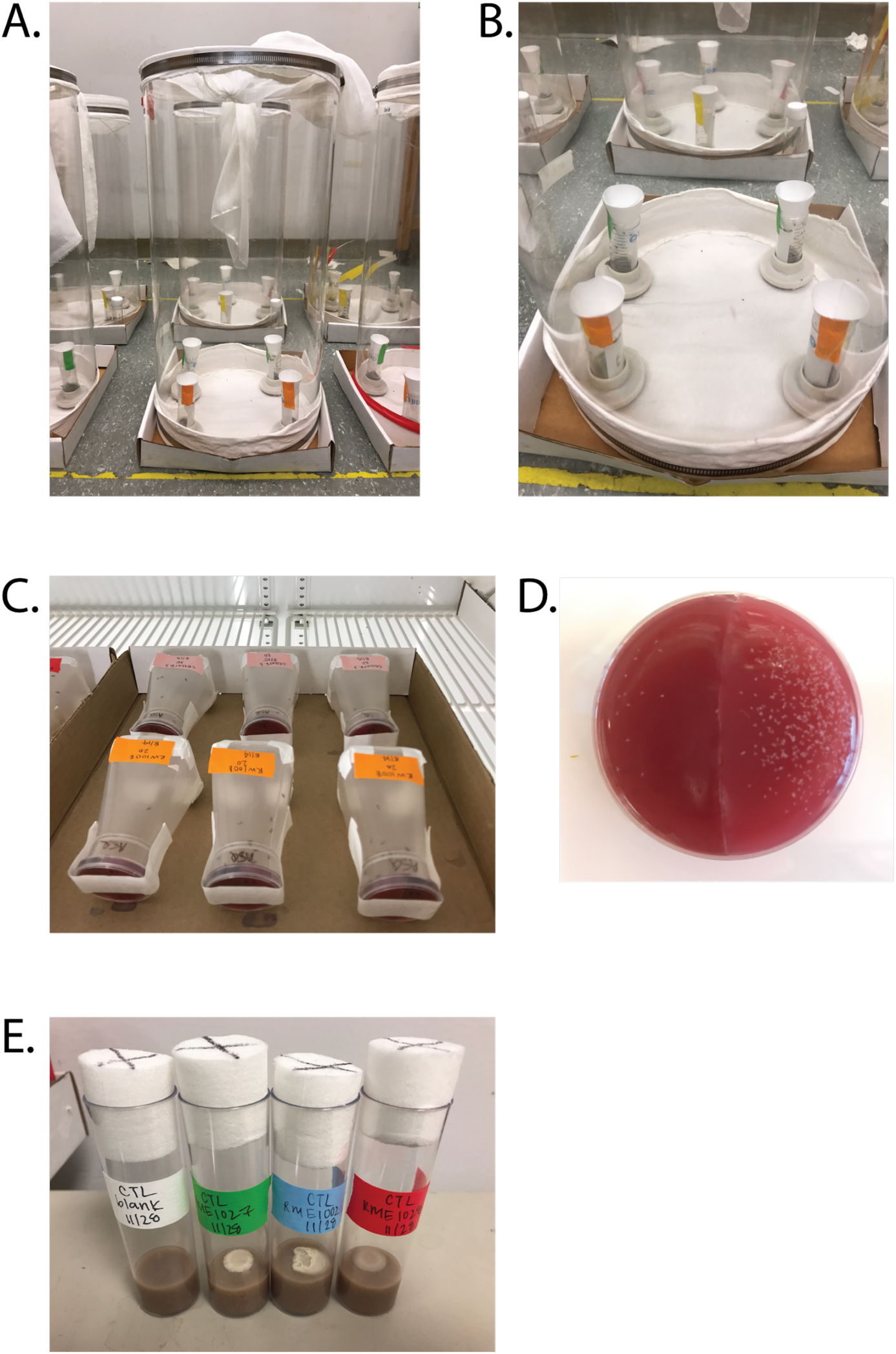
Photographs of behavior assays. (A) Setup of a single, trap-based olfactory assay. (B) Close up of trap-based olfactory assay. Traps can be arranged in four possible combinations, two of which are depicted here. (C) Setup of six oviposition assays. (D) Example agar plate after oviposition assay. Left side is uninnoculated grape juice agar, right side is yeast inoculated grape juice agar. (E) Larval development and longevity assays were performed in wide vials shown here. Both larvae and adults were exposed to a live monoculture of yeast spotted onto sterile banana media. Depicted are Day 7, negative control vials that had no larvae or adult flies but grew alongside behavioral assays to monitor bacterial or mold contamination.

**Figure S6.**
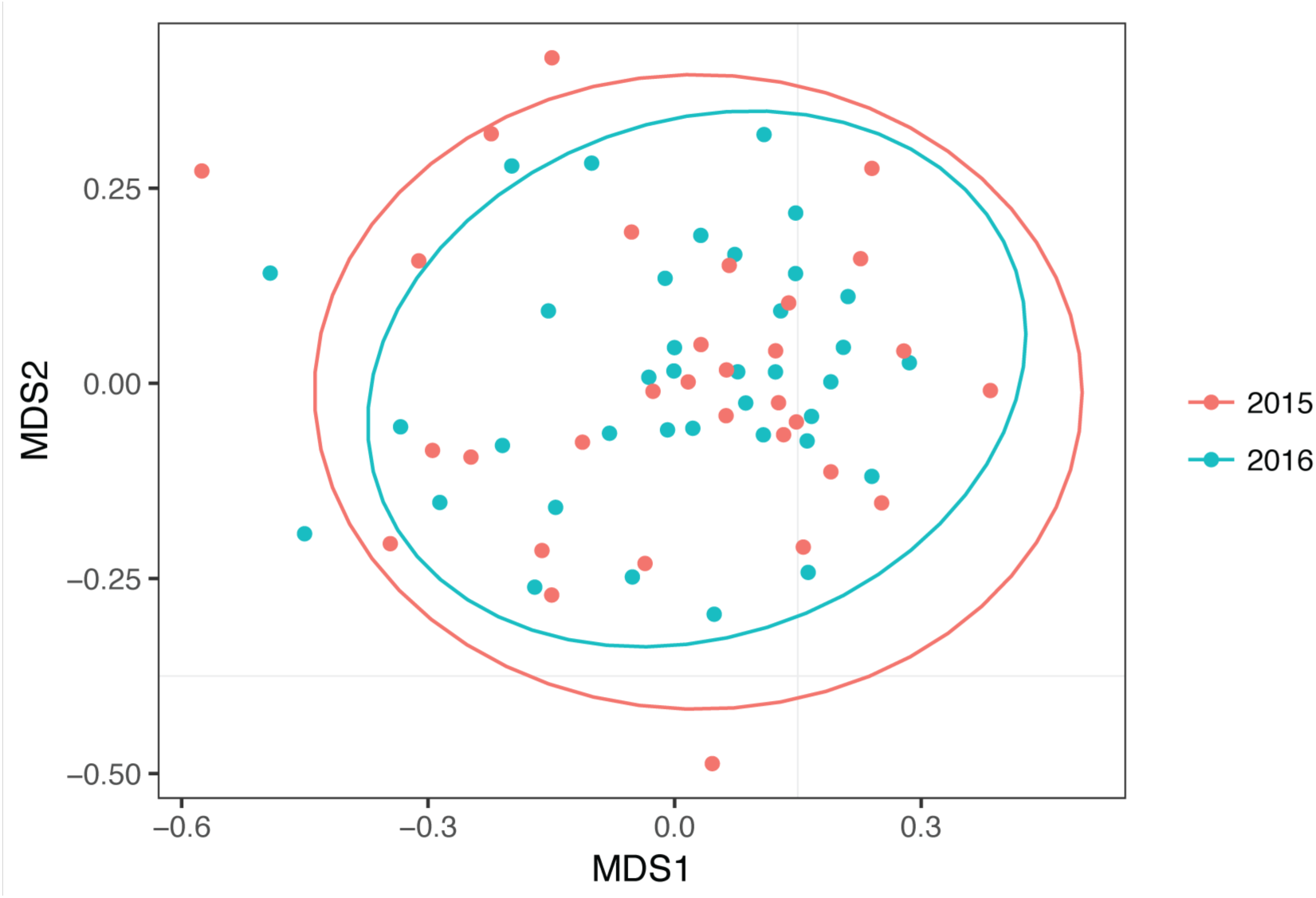
Bray-Curtis dissimilarity NMDS of PCR plate sequenced with samples collected in 2015 at UC Berkeley and the same plate resequenced with samples collected in 2016 at UC Davis.Each sample was rarefied to 200 sequences and is represented by a single point, color-coded by year.

**Table S1.** Library and quality filtering statistics

|  | 2015 total<br>reads | 2016 total<br>reads | Total<br>OTUs | % of original<br>reads<br>retained |
| --- | --- | --- | --- | --- |
| Raw sequences | 8724595 | 15570284 | -- | -- |
| After merging pairs | 8034126 | 14261774 | -- | 91.77% |
| After demultiplexing | 5389671 | 7311698 | -- | 52.28% |
|  | <u>Combined total reads</u> |  |  |  |
| Removal of resequenced control | 12037090 |  | -- | 49.55% |
| After quality filtering | 11302418 |  | -- | 46.52% |
| After chimera filtering | 11293777 |  | 6159 | 46.49% |
| Post OTU picking quality filtering | 7609820 |  | 399 | 31.32% |
| Final sequence count | 7609820 |  | 399 | 31.32% |

**Table S2.**
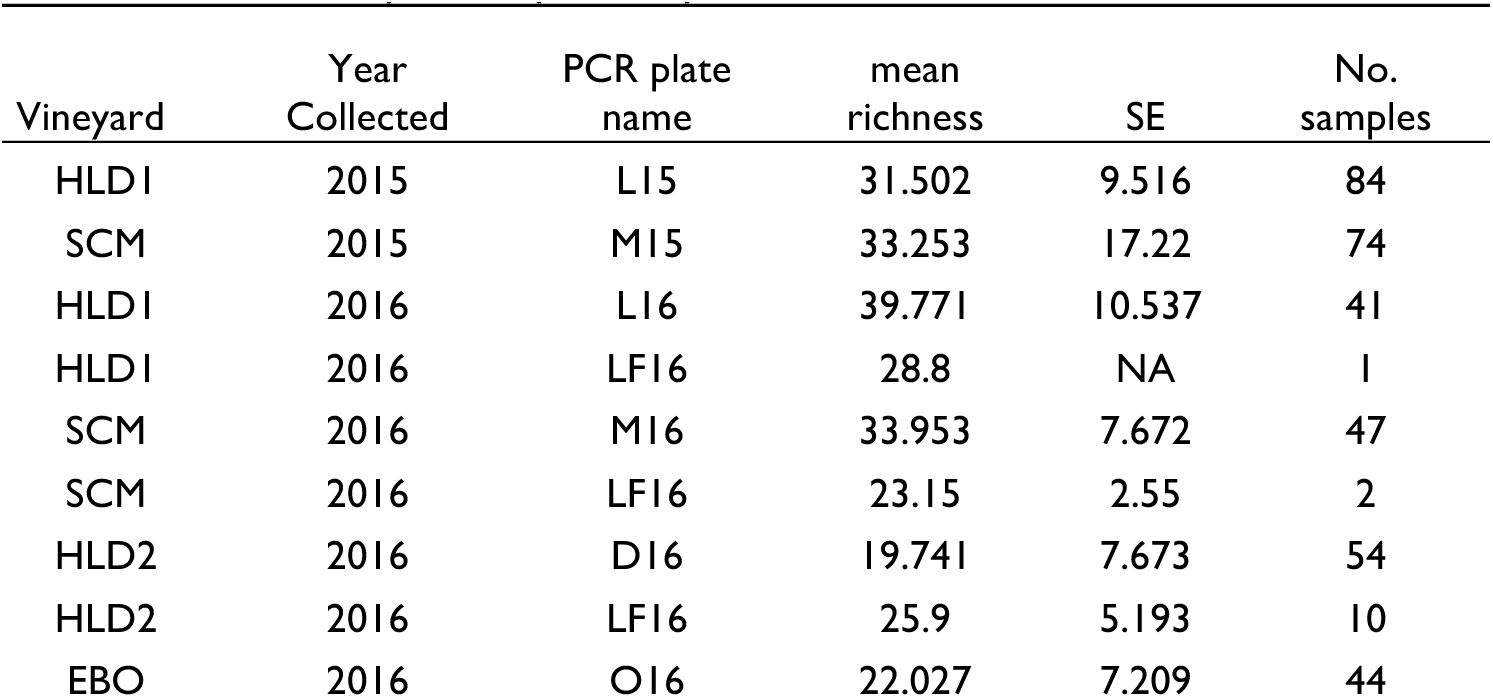
Alpha richness of fungi vectored by *Drosophila* by vineyard and harvest season, rarefied to 1000 sequences per sample.

**Table S3.**
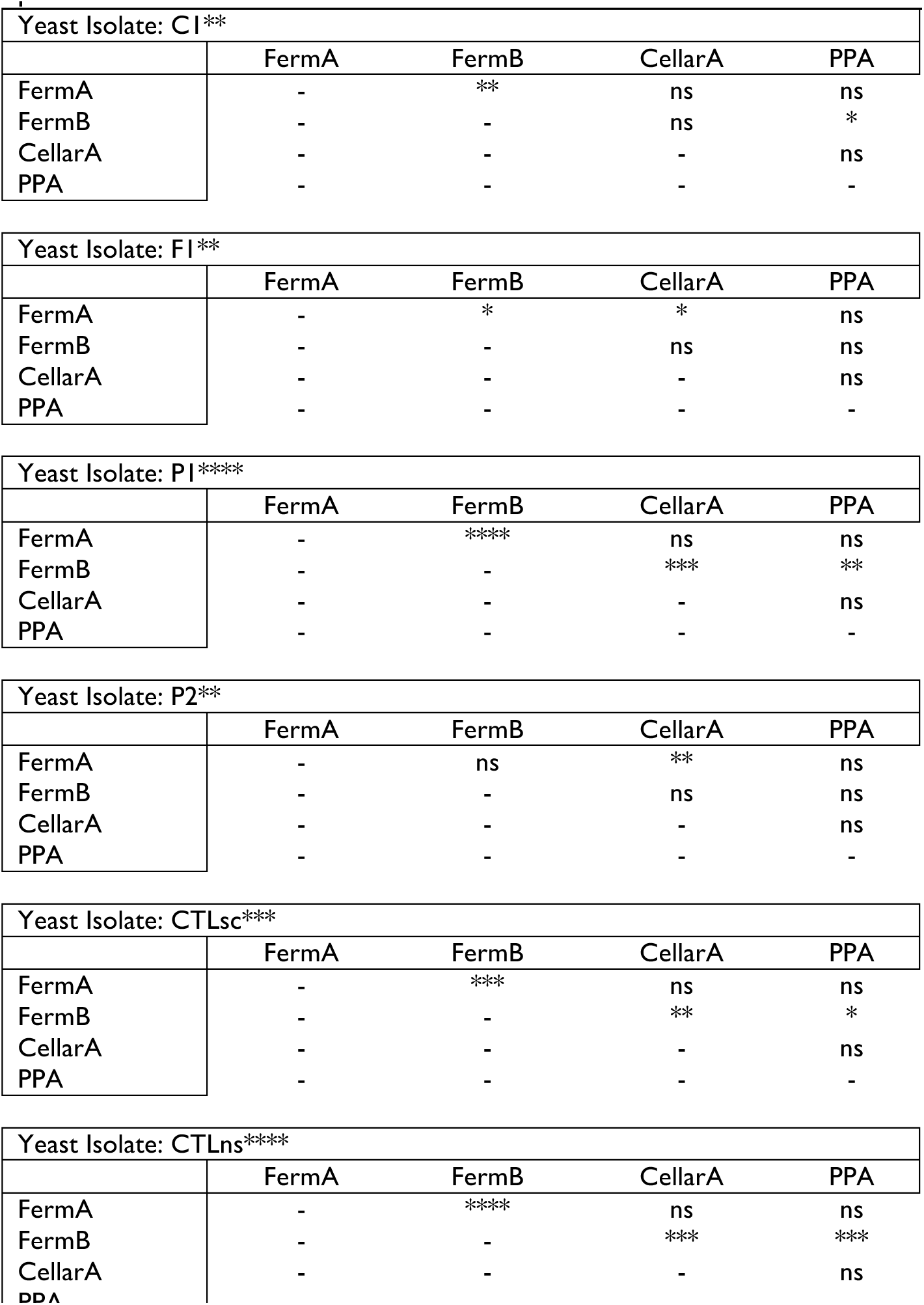
Statistics accompanying the ovipostion index responses of each fly line in Figure 4 for each yeast species. Each table represents a different yeast species. Within each yeast species, ANOVA was first used to test for any differences in ovipostion responses between fly lines for a given yeast species and denoted by a * next to each yeast isolate. If ANOVA results were statistically significant, Tukey’s multiple comparisons test was used to identify the fly lines exhibiting ovipostion responses that were significantly different than other lines and are depicted with * within the table. ns: not significant, *: p<0.05, **: p<0.01, ***: p<0.001, ****: p<0.0001.

**Table S4.** Average percentage of *Drosophila* larvae that eclose successfully when developing on different yeast species. Control conditions are shaded grey. (ANOVA when compared to no yeast control followed by Tukey’s multiple comparisons test was used to calculate significance values, ****:p < 0.0001).

| Fly Line | ANOVA | Yeast treatment |  |  |  |  |  | no yeast | Dead yeast, cornmeal media |
| --- | --- | --- | --- | --- | --- | --- | --- | --- | --- |
|  |  | CI | FI | PI | P2 | CTLsc | CTLns |  |  |
| FermA | **** | 88.9% | 100.0% | 100.0% | 100.0% | 100.0% | 97.8% | 0.0% | 94.4% |
| FermB | **** | 88.9% | 100.0% | 93.3% | 82.2% | 88.9% | 100.0% | 0.0% | 93.3% |
| CellarA | **** | 64.4% | 100.0% | 91.1% | 93.3% | 88.9% | 95.6% | 0.0% | 96.7% |
| PPA | **** | 80.0% | 91.1% | 91.1% | 93.3% | 86.7% | 84.4% | 20.0% | 67.8% |

**Table S5.** Statistics to accompany Table 2 on effects of yeast species on larval development time for each fly line. (One way ANOVA followed by Tukey’s multiple comparisons test was used to calculate significance values, ****: p<l0.0001).

| **FermA** | CI | FI | PI | P2 | CTLsc | CTLns |
| --- | --- | --- | --- | --- | --- | --- |
| CI | - | **** | **** | **** | **** | **** |
| FI | - | - | ns | ns | ns | ns |
| PI | - | - | - | ns | ns | ns |
| P2 | - | - | - | - | ns | ns |
| CTLsc | - | - | - | - | - | ns |
| CTLns | - | - | - | - | - | - |

| **FermB** | CI | FI | PI | P2 | CTLsc | CTLns |
| --- | --- | --- | --- | --- | --- | --- |
| CI | - | **** | **** | **** | **** | **** |
| FI | - | - | ns | ns | ns | ns |
| PI | - | - | - | ns | ns | ns |
| P2 | - | - | - | - | ns | ns |
| CTLsc | - | - | - | - | - | ns |
| CTLns | - | - | - | - | - | - |

| **CellarA** | CI | FI | PI | P2 | CTLsc | CTLns |
| --- | --- | --- | --- | --- | --- | --- |
| CI | - | **** | **** | **** | **** | **** |
| FI | - | - | ns | ns | ns | ns |
| PI | - | - | - | ns | ns | ns |
| P2 | - | - | - | - | ns | ns |
| CTLsc | - | - | - | - | - | ns |
| CTLns | - | - | - | - | - | - |

| **PPA** | CI | FI | PI | P2 | CTLsc | CTLns |
| --- | --- | --- | --- | --- | --- | --- |
| CI | - | **** | **** | **** | **** | **** |
| FI | - | - | ns | ns | ns | ns |
| PI | - | - | - | ns | ns | ns |
| P2 | - | - | - | - | ns | ns |
| CTLsc | - | - | - | - | - | ns |
| CTLns | - | - | - | - | - | - |

**Table S6.** Comparisons of fly line lifespans to each other. One way ANOVA followed by Tukey’s multiple comparisons test was used to calculate significance values, ns: not significant,*:p<0.05,**:p<0.01,***:p<0.001,****:p<0.0001.

|  | FermA | FermB | CellarA | PPA |
| --- | --- | --- | --- | --- |
| FermA | - | * | * | ns |
| FermB | - | - | ns | *** |
| CellarA | - | - | - | ** |
| PPA | - | - | - | - |

**Table S7.** Media and buffers used in this study.

| Media name | Purpose | Volume | Recipe |
| --- | --- | --- | --- |
| 5% YPD, agar | Solid media for yeast isolation | 1L | 20g Peptone (BD Bacto Peptone), 10g Yeast Extract (Amresco Yeast Extract, Bacteriological, Ultra Pure Grade), 50g Dextrose (Fisher Scientific Dextrose Anhydrous), 20g Agar (BD Bacto Agar), MilliQ water to 1L. Autoclaved and poured into 100 x 60mm petri dishes (Falcon). |
| WLB | Buffer for DNA extration | - | 2M Guanidinium thiocyanate (Fisher Scientific), 0.5 M EDTA (Fisher Scientific), 1.8% Tris base (Promega), 8% NaCl (Sigma Aldrich), 150mL of MilliQ water, and adjust to pH 8.5. Autoclaved and filter sterilized (Nalgene 75mm filter unit, 0.2aPES). (Will Ludington, personal communication). |
| Agar grape juice | Olfactory preference assay | 1L | 1.7g Yeast nitrogen base without amino acids or ammonium sulfate (Difco, BD), 20g Agar (BD Bacto Agar), 355mL Organic Cascadian Farms Concord grape juice concentrate (1 can), and 645mL MilliQ water. Heated to a boil and poured into 60 x 15mm petri dishes (Falcon). |
| Liquid grape juice | Oviposition assay | 1L | (Prepare as instructed on can) 1 can of Organic Cascadian Farms Concord grape juice concentrate, 3 parts MilliQ water. Heated to a boil. |
| 5% YPD, liquid | Liquid media for yeast starter cultures | 1L | 20g Peptone (BD Bacto Peptone), 10g Yeast Extract (Amresco Yeast Extract, Bacteriological, Ultra Pure Grade), 50g Dextrose (Fisher Scientific Dextrose Anhydrous), and MilliQ water to 1L. Autoclaved and filter sterilized (Nalgene 75mm filter unit, 0.2aPES). |
| GB media | Larval development and longevity assay | see recipe | 40% weight by volume (w/v) fresh, pureed organic banana, 60% w/v MilliQ water, and 1.25% agar (BD Bacto Agar). Autoclaved and poured into wide mouth *Drosophila* vials (wide mouth, K-resin, Genessee Scientific). |

**Table S8.** Components of mock community and sequencing results. Mock culture replicates were independent with the exception of “2016 reseq of 2015” which came from amplicons made in 2015 and resequenced in 2016 as a control. Yeast isolates used in mock cultures were isolated, species were identified, and DNA was extracted as described in Materials and Methods. The mock culture was created by combining 4uL of each yeast species listed below. The resulting mix was used as DNA template in the amplification protocol described.

| Yeast species | Conc.<br>(ng/uL) | 2015 MC<br>Rep#1,<br>ratio of<br>sequences | 2015 MC<br>Rep#2,<br>ratio of<br>sequences | 2015 MC<br>Rep#3,<br>ratio of<br>sequences | 2016 reseq<br>of 2015<br>MC<br>Rep#1,<br>ratio of<br>sequences | 2016 reseq<br>of 2015<br>MC<br>Rep#2,<br>ratio of<br>sequences | 2016 reseq<br>of 2015<br>MC<br>Rep#3,<br>ratio of<br>sequences | 2016 MC<br>Rep#1,<br>ratio of<br>sequences | 2016 MC<br>Rep#2,<br>ratio of<br>sequences | 2016 MC<br>Rep#3,<br>ratio of<br>sequences |
| --- | --- | --- | --- | --- | --- | --- | --- | --- | --- | --- |
| *Candida parapsilosis* | 21.75 | 0.22 | 0.22 | 0.22 | 0.22 | 0.22 | 0.22 | 0.20 | 0.22 | 0.24 |
| *Lodderomyces elongisporus* | 15.1 | 0.18 | 0.19 | 0.19 | 0.19 | 0.20 | 0.19 | 0.15 | 0.16 | 0.17 |
| *Pichia fermentans* | 1.24 | 0.00 | 0.00 | 0.00 | 0.00 | 0.00 | 0.00 | 0.00 | 0.00 | 0.00 |
| *Candida boidinii* | 15.6 | 0.35 | 0.33 | 0.35 | 0.39 | 0.39 | 0.36 | 0.13 | 0.26 | 0.29 |
| *Saccharomyces cerevisiae* | 4.32 | 0.02 | 0.02 | 0.02 | 0.02 | 0.02 | 0.02 | 0.05 | 0.07 | 0.09 |
| *Hanseniaspora uvarum* | 14 | 0.01 | 0.02 | 0.01 | 0.01 | 0.01 | 0.02 | 0.21 | 0.08 | 0.04 |
| *Issachenkia terricola* | 16.25 | 0.21 | 0.23 | 0.20 | 0.17 | 0.15 | 0.18 | 0.26 | 0.21 | 0.18 |
| *Metschnikowia pulcherrima* | 11.55 | 0.00 | 0.00 | 0.00 | 0.00 | 0.00 | 0.00 | 0.00 | 0.00 | 0.00 |
| *Zygoascus meyeriae* | 20.2 | 0.00 | 0.00 | 0.00 | 0.00 | 0.00 | 0.00 | 0.00 | 0.00 | 0.00 |

